# Improved constructs for bait RNA display in a bacterial three-hybrid assay

**DOI:** 10.1101/2024.07.23.604302

**Authors:** Linh D. Nguyen, Hannah LeBlanc, Katherine E. Berry

## Abstract

We have previously developed a transcription-based bacterial three-hybrid (B3H) assay as a genetic approach to probe RNA-protein interactions inside of *E. coli* cells. This system offers a straightforward path to identify and assess the consequences of mutations in RBPs with molecular phenotypes of interest. One limiting factor in detecting RNA-protein interactions in the B3H assay is RNA misfolding arising from incorrect base-pair interactions with neighboring RNA sequences in a hybrid RNA. To support correct folding of hybrid bait RNAs, we have explored the use of a highly stable stem (“GC clamp”) to isolate regions of a hybrid RNA as discrete folding units. In this work, we introduce new bait RNA constructs to 1) insulate the folding of individual components of the hybrid RNA with GC clamps and 2) express bait RNAs that do not encode their own intrinsic terminator. We find that short GC clamps (5 or 7 bp long) are more effective than a longer 13bp GC clamp in the B3H assay. These new constructs increase the number of Hfq-sRNA and -5′UTR interactions that are detectable in the B3H system and improve the signal-to-noise ratio of many of these interactions. We therefore recommend the use of constructs containing short GC clamps for the expression of future B3H bait RNAs. With these new constructs, a broader range of RNA-protein interactions are detectable in the B3H assay, expanding the utility and impact of this genetic tool as a platform to search for and interrogate mechanisms of additional RNA-protein interactions.

## INTRODUCTION

RNA-protein interactions play essential roles in regulating gene expression in all organisms. In bacteria, several classes of global RNA-binding proteins (RBPs) are known to contribute to the stability and interactions of small RNAs (sRNAs) and the translation and turnover of messenger RNAs (mRNAs) (Holmqvist et al. 2020; Holmqvist and Vogel 2018; Woodson et al. 2018; Gottesman and Storz 2015; Olejniczak and Storz 2017; Gottesman and Storz 2015). For instance, the paradigmatic bacterial RNA-chaperone protein Hfq utilizes distinct faces of its toroid surface to mediate specific interactions between sRNAs and mRNA 5′ untranslated regions (5′UTRs) to regulate recruitment of the ribosome or RNA degradosome (Updegrove et al. 2016; Woodson et al. 2018; Kavita et al. 2018; Wagner and Romby 2015). Despite substantial progress in understanding the details of Hfq-facilitated sRNA-based gene regulation, significant questions remain about the molecular mechanisms other global RBPs use to interact with RNA, how the structure and function of these proteins vary across bacterial species, and what additional as-yet-undiscovered RBPs may support the function of ncRNAs in bacterial species without global RBPs such as ProQ and Hfq.

To improve our understanding of the scope and mechanisms of bacterial RBPs, we have developed a genetic assay to assess the strength of RNA-protein interactions in a living *E. coli* reporter cell (Berry and Hochschild 2018; Stockert et al. 2022). This bacterial three-hybrid (B3H) assay utilizes *E. coli*’s cellular machinery to express hybrid RNAs and proteins *in vivo*, avoiding the need to purify these components and allowing them to be studied in native cytoplasmic conditions. We have used the B3H system to study the interactions of both *E. coli* Hfq and ProQ (Berry and Hochschild 2018; Pandey et al. 2020; Wang et al. 2021; Stein et al. 2020, 2023). As the assay affords access to powerful tools of molecular genetics, it supports the dissection of molecular mechanisms using unbiased mutagenesis screens, as well as the potential discovery of novel RBPs.

While there is substantial potential in the B3H assay, its current utility has been limited by only a modest success rate in detecting established RNA-protein interactions. Improvements in the expression level of adapter protein have increased signal for several Hfq-sRNA interactions with K_D_ values out to ∼100 nM (Wang et al. 2021). Despite this progress, many well-characterized Hfq-sRNA interactions are undetectable in the current B3H assay. In addition, the B3H assay has only been used to detect interactions with RNAs that encode their own intrinsic terminators, even though interactions with 5′UTRs are known to play important regulatory roles in cells, and fragments of many open reading frames (ORFs) have been identified in transcriptome-wide RNA binding studies of global RBPs such as Hfq, ProQ and KH-domain proteins (Melamed et al. 2020; Holmqvist et al. 2016, 2018; Zhu et al. 2024; Lamm-Schmidt et al. 2021; Michaux et al. 2023; Smirnov et al. 2016; Hör et al. 2020a).

One constraint on the B3H assay is that an RNA of interest must be expressed as a hybrid RNA tethered to other elements such as an MS2 hairpin (MS2^hp^). Due to shallow folding energy landscapes, hybrid RNA sequences have high potential to misfold. Prior work in an analogous yeast three-hybrid (Y3H) assay has demonstrated that GC clamps can be useful in isolating discrete units of RNA secondary structure and facilitating the correct folding of hybrid RNAs (Cassiday and Maher 2001; Bernstein 2002). Based on this precedent, we have developed and tested updated constructs for the display of bait RNAs — so called “pBait” plasmids — in our B3H assay. In addition to optimizing constructs for the display of RNAs that encode their own intrinsic terminator (*e*.*g*. sRNAs and 3′UTRs), we have also developed constructs that provide an exogenous intrinsic terminator for use with RNA sequences that do not contain a terminator, *e*.*g*. 5′UTRs, ORF fragments, or non-bacterial RNAs. In this report, we demonstrate that the presence of a GC clamp in bait RNA constructs improves the signal-to-noise ratio of many B3H interactions — for both RNAs that do and do not encode their own intrinsic terminator. In both cases, a shorter GC clamp than what was previously used in the Y3H assay is most successful in the bacterial system. Based on these findings, we recommend using pBait constructs with GC clamps that are 5-to-7 base pairs long to clone future RNA baits for the B3H assay.

## RESULTS AND DISCUSSION

### Introduction to bacterial three-hybrid assay

The bacterial three-hybrid (B3H) assay is a useful tool for detection and characterization of RNA-protein interactions in *E. coli* (Berry and Hochschild 2018, Stockert et al. 2022) and has been used to investigate interactions of sRNAs and 3′UTRs with the RBPs Hfq and ProQ (Berry and Hochschild 2018, Wang et al. 2021, Pandey et al. 2020, Stein et al. 2020, 2023). Fundamentally, the B3H assay detects RNA-protein interactions by connecting the strength of the interaction to the expression of a reporter gene *lacZ*, through interaction of three hybrid components: a “bait” RNA, a “prey” protein, and an RNA-DNA adapter (Fig. 1A). RNAs of interest can be inserted in between XmaI and HindIII restriction sites on a “pBait” plasmid to express a hybrid RNA with the bait RNA downstream of an MS2 hairpin (MS2^hp^) (Fig. 1B). The interaction between “bait” RNA tethered upstream of the promoter through the MS2^hp^ and the “prey” protein — fused to the α subunit of RNA polymerase (RNAP) and expressed from a “pPrey” plasmid — stabilizes RNAP at the promoter and increases transcription of *lacZ*. The interaction can be quantified as the fold-stimulation of β-galactosidase (β-gal) activity. For example, when pBait-ChiX and pPrey-Hfq are transformed into reporter cells along with pAdapter, β-gal activity is stimulated ∼4.5-fold above the background activity when either prey, bait or adapter is missing from the transformed plasmids (Fig. 1C). In practice, we have found that B3H interactions above 2-fold stimulation tend to be robust, repeatable and useful for both forward and reverse mutagenic analysis (Pandey et al. 2020; Berry and Hochschild 2018; Stockert et al. 2022).

**Figure 1.**
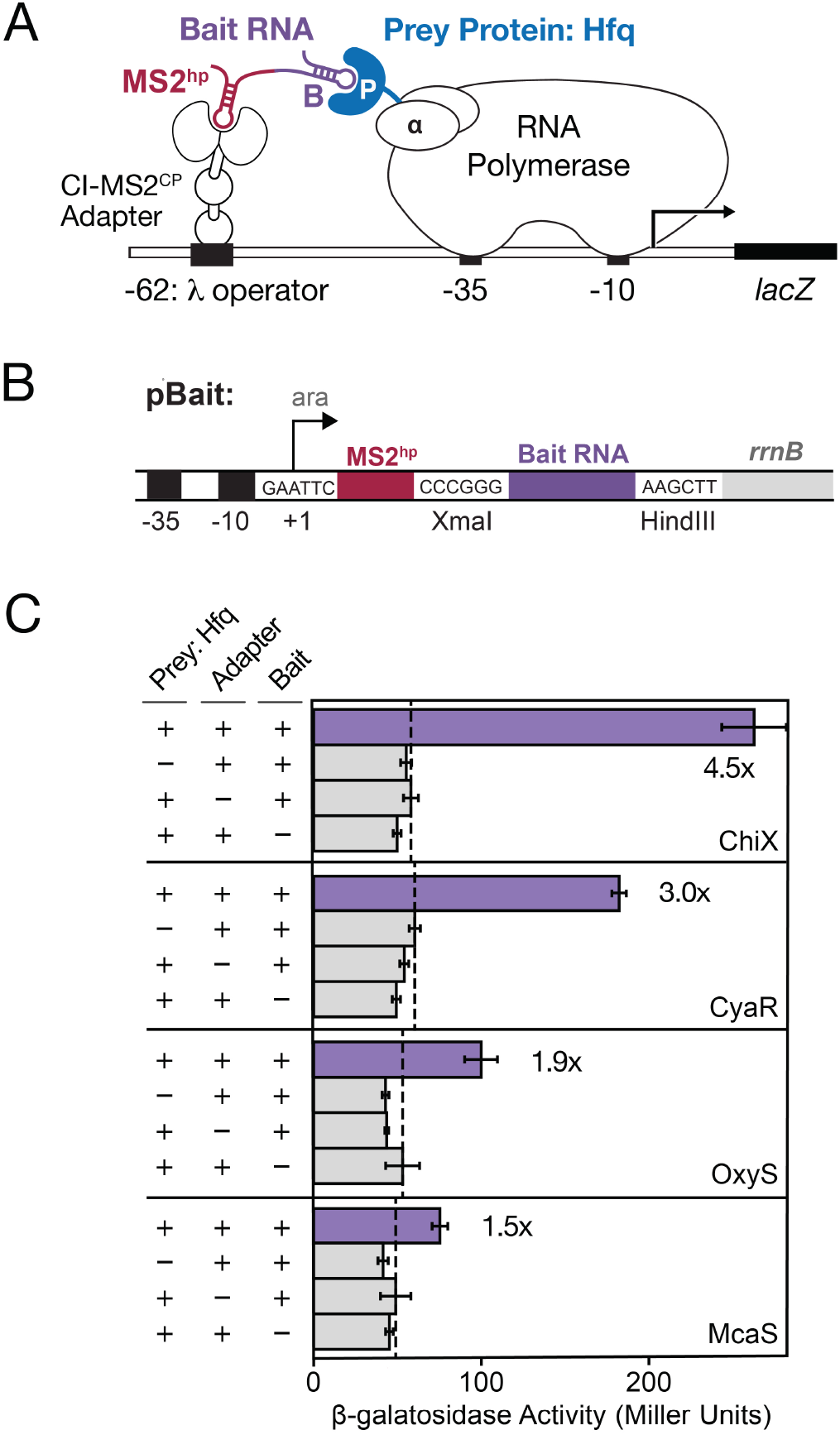
Bacterial three-hybrid (B3H) assay enables detection of a subset of Hfq-sRNA interactions. (A) Schematic showing B3H system for detection of Hfq-RNA interactions. Interaction between an Hfq “prey” protein moiety and RNA “bait” moiety activates transcription from a test promoter, which directs transcription of a *lacZ* reporter gene. The test promoter (p*lac*-O_L_2–62), bears the λ operator O_L_2 centered at position –62 relative to the transcription start site. The RNA-binding moiety MS2^CP^ is fused to λCI (CI-MS2^CP^) as an RNA-DNA adapter (“Adapter”) to tether the hybrid bait RNA (MS2^hp^-B) to the test promoter. The prey protein is C-terminally fused to the NTD of the α subunit of RNA polymerase. (B) Schematic showing 1xMS2^hp^ pBait construct with XmaI and HindIII restriction sites present for insertion of bait RNAs. The likely transcription start site (the second A within an EcoRI site upstream of the MS2^hp^) is indicated. (C) Results of β-galactosidase (β-gal) assays performed in KB483 cells transformed with three compatible plasmids: pPrey that encoded α alone (–) or the α-Hfq fusion protein (+; pKB817), pAdapter that encoded λCI alone (–) or the CI-MS2^CP^ fusion protein (+; p35u4), and pBait that encoded a hybrid RNA with either the ChiX, CyaR, OxyS or McaS sequence sequence following one copy of an MS2^hp^ moiety (+; pCH6, pCH8, pCH9 or pCH7, respectively) or an RNA that contained only the MS2^hp^ moiety (–). Fold-stimulation over basal levels are indicated: that is, the β-gal activity measured in the presence of all hybrid constructs divided by the activity of the highest negative control sample expressing only part of one hybrid construct. The dashed vertical lines represent “basal” levels.

While the original pBait constructs contained two copies of the MS2^hp^ (Berry and Hochschild 2018; Wang et al. 2021), a single MS2^hp^ has been shown to be sufficient for detection of Hfq-sRNA interactions and, indeed, is critical for accurate detection of ProQ-RNA interactions (Pandey et al. 2020). Due to its simplicity and broader applicability, we chose to focus our optimization of RNA display here on the 1xMS2^hp^ pBait construct. With the 1xMS2^hp^ pBait construct, Hfq interactions with sRNAs such as CyaR and ChiX are detected at 3-to 5-fold stimulation of *lacZ* activity (Fig. 1C), but interactions with sRNAs such as McaS and OxyS are more weakly detected (1.5-to 2-fold stimulation). To maximize the utility of the bacterial three-hybrid assay, a larger fraction of known RNA-protein interactions and classes must be successfully and robustly detectable.

### Predicted folding of hybrid RNAs

Considering factors that could limit the detection of certain Hfq-sRNA interactions, we explored the possibility that hybrid RNAs could misfold due to the additional elements present in the hybrid RNA construct (Fig. 1A,B). Using an RNA secondary structure visualization tool called forna (Kerpedjiev et al. 2015), we found that many “bait” RNAs were indeed predicted to misfold when fused to an MS2^hp^. For instance, when the sRNA McaS is cloned into the original 1xMS2^hp^ pBait construct (Fig. 2A), it is predicted to fold quite differently than when expressed alone (Fig. 2B). The same was true to varying extents for hybrid RNAs containing CyaR, OxyS and ChiX (Supplemental Fig. S1). Such misfolding of an sRNA would likely disrupt its interaction with the Hfq “prey” protein, and could be a limiting factor in the B3H assay’s detection of RNA-protein interactions.

**Figure 2.**
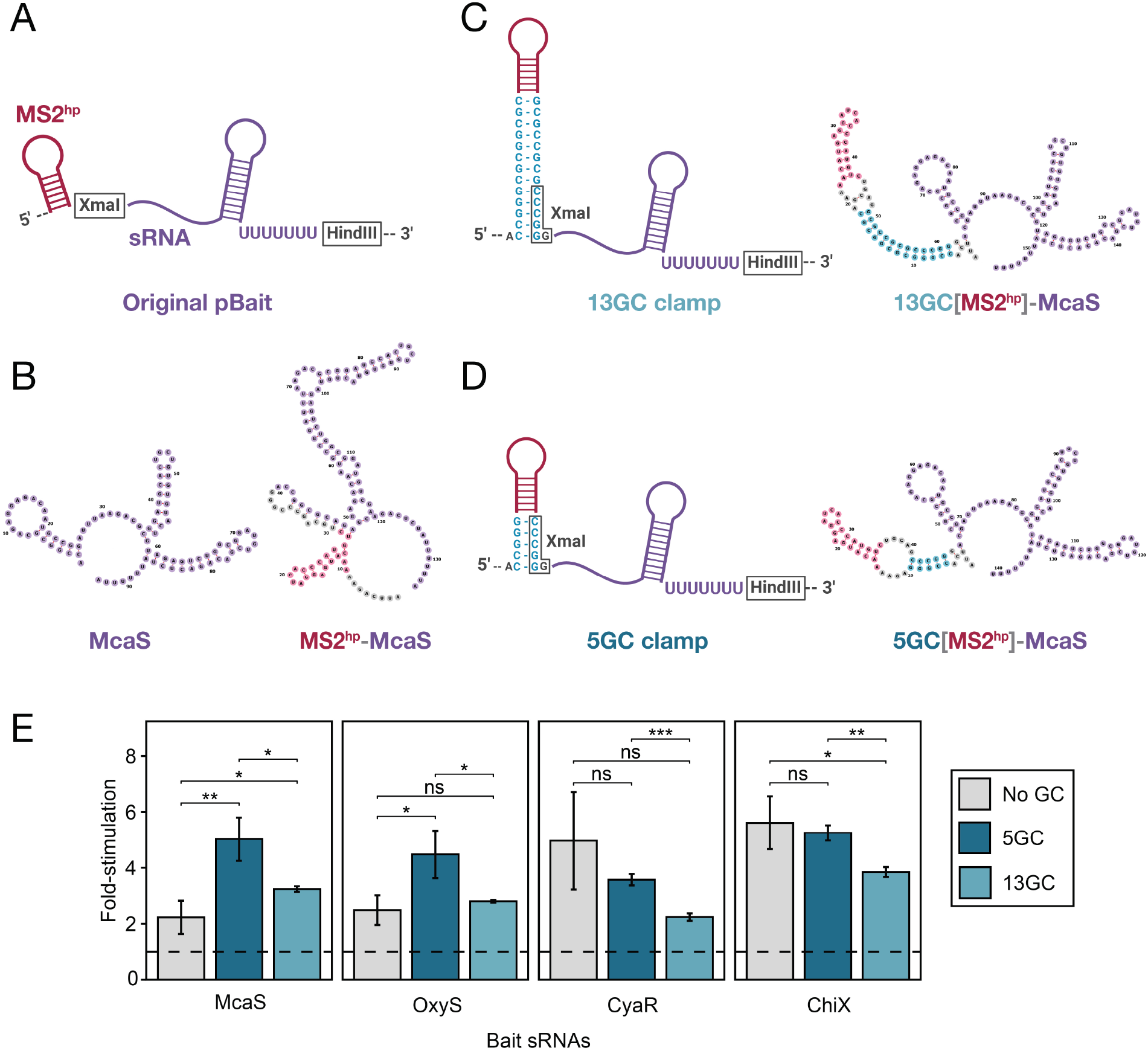
A short GC clamp is more effective than a longer clamp in improving B3H signal for Hfq-sRNA interactions. (A) Schematic of original hybrid bait RNA construct showing the intended secondary structure of the MS2^hp^ and an sRNA inserted between the XmaI and HindIII restriction sites, expressing its own intrinsic terminator. (B) forna predictions of the *E. coli* McaS sRNA on its own (left) and when expressed as a hybrid RNA in the original pBait plasmid (right). (C,D) Schematic of hybrid RNAs with additional GC clamps that are 13bp (C) or 5bp (D) long around the MS2^hp^. Schematics show intended secondary structure (left) and forna-predicted secondary structures of McaS expressed in these hybrid RNAs (right). (e) Results of β-gal assays performed in KB483 cells transformed with three compatible plasmids: pPrey that encoded α alone (–) or the α-Hfq fusion protein (pKB817), pAdapter that encoded λCI alone (–) or the CI-MS2^CP^ fusion protein (p35u4), and pBait that encoded a hybrid RNA with either the ChiX, CyaR, OxyS or McaS sequence sequence following one copy of an MS2^hp^ moiety in the absence of a GC clamp (“No GC”; pCH6, pCH8, pCH9 or pCH7, respectively), or the presence of a GC clamp flanking the MS2^hp^ moiety of either 5bp (“5GC”; pLN28, pLN35, pLN36 or pLN34, respectively) or 13bp (“13GC”; pLN27, pLN32, pLN33 or pLN31, respectively), or an RNA that contained only the MS2^hp^ moiety (pCH1, pSS1 or pSW1). Bar graphs here and throughout the rest of the paper show B3H interactions as fold-stimulation over basal levels, indicated by a dashed horizontal line. Data are presented as the average value of three measurements taken on multiple days and error bars show the standard deviation of these values. Student t-tests were conducted to assess statistical significance for differences between pairs of B3H interactions; asterisks indicate P-values, represented as follows: ns, not significant, ^*^P < 0.05, ^**^P < 0.01, ^***^P< 0.001).

Previous studies of RNA-protein interactions using a yeast three-hybrid (Y3H) assay demonstrated that the introduction of a “GC clamp” flanking an RNA-of-interest improved detection of RNA-protein interactions of the p50 protein (Cassiday and Maher 2001). The GC clamp used in the Y3H assay is a 13-bp stretch of complementary guanosine (G) and cytidine (C) residues (Cassiday and Maher 2001; Bernstein 2002). Since G-C base pairs are more stable than A-U or G-U pairing, complementary strands in the “clamp” are more likely to base pair with one another than unintended sequences elsewhere in the RNA; this can help insulate secondary structure elements within the hybrid RNA.

To explore whether the presence of a GC clamp would support accurate folding of hybrid bait RNA constructs in the B3H assay, we examined the predicted secondary structures of RNAs in the presence of the GC clamps using forna (Kerpedjiev et al. 2015). We found that a 13-bp GC clamp on either side of the MS2^hp^ was predicted to promote proper folding of hybrid RNAs containing the McaS sRNA (Fig. 2C) as well as OxyS, CyaR and ChiX (Supplemental Fig. S1). Because the dsRNA in a 13-bp GC clamp has the potential to be structurally quite rigid, we wondered whether a shorter GC clamp might provide more flexibility in the orientation of the hybrid RNA and favor formation of bait-prey interactions. We therefore tested the effects of shorter GC clamps on the predicted folding of bait RNAs. Strikingly, shorter GC clamps — down to 5 bp — were still predicted to facilitate the proper folding of all sRNAs (Fig. 2D and Supplemental Fig. S1). To increase the chances of improving the B3H signal with new pBait constructs, we chose to compare the effects of two GC clamp constructs: one with the original length used in the Y3H assay (“long”; 13 bp; Fig. 2C) and one with the minimum length predicted to support correct folding (“short”; 5 bp; Fig. 2D).

### A short GC clamp improves B3H signal for sRNAs

To determine the effects of long and short GC-clamp modifications in B3H pBait constructs, we tested the interactions of several Hfq-dependent *E. coli* sRNAs (McaS, CyaR, OxyS and ChiX) with Hfq in the original pBait construct vs. the two new GC-clamp-containing constructs. In the presence of a short GC clamp (“5GC”), the signal for interactions of McaS and OxyS with Hfq increases in a statistically significant manner, from just over 2-fold to greater than 4-fold stimulation of *lacZ* activity (Fig. 2E). In contrast, the interactions that were already stronger than 4-fold stimulation (CyaR-Hfq and ChiX-Hfq) were not significantly altered by this modification. As hypothesized, the long GC clamp (“13GC”) was not as effective as the short clamp in improving B3H signal. Only one hybrid RNA (McaS) showed a statistically significant increase in signal with the 13GC clamp compared to the original construct, and all four hybrid RNAs demonstrated higher signal with the shorter clamp than the longer one (Fig. 2E). Together, these results suggest that the short GC clamp is a promising construct to improve the detection of sRNA-Hfq interactions in the B3H assay. This effect is likely mediated by facilitating the correct folding of bait RNAs and MS2^hp^ in the context of the hybrid RNA.

### Expansion of detectable interactions between Hfq and sRNA targets

Having demonstrated the potential of a short GC clamp to improve the detection of several sRNA-Hfq interactions, we wanted to explore the effects of this construct on a broader set of interactions. We selected a group of *E. coli* sRNAs that had been found to interact with Hfq *in vivo* (Zhang et al. 2003; Melamed et al. 2020; Hör et al. 2020b), including some whose B3H interactions with Hfq were previously found to be low (ArcZ, MicF, DsrA) or undetectable (RyhB, SgrS, GcvB, GlmZ, Spot42) (Berry and Hochschild 2018; Wang et al. 2021), as well as some that had not been previously tested in the B3H assay (DicF, OmrA, SdhX, MicL, GadY). All sRNAs were cloned into the 5GC[1xMS2^hp^] pBait construct, and newly cloned sRNAs were also inserted into the 1xMS2^hp^ pBait construct (Fig. 2A). For the subset of the sRNAs that had been previously tested in 2xMS2^hp^ constructs (Spot42, MicF, GcvB, DsrA, GlmZ), the interactions of the new 5GC[1xMS2^hp^] constructs were compared to the original 2xMS2^hp^ constructs which had produced low or undetectable interactions with Hfq (Supplemental Figs. S2 and S3). In B3H experiments comparing the strength of sRNA-Hfq interactions with new vs. old constructs, the 5GC pBait construct statistically significantly improved 8 of the 15 tested sRNA-Hfq B3H interactions, compared to the original pBait construct (Fig. 3). The short GC clamp did not produce significant changes in B3H signal for another 6 of the 15 sRNAs, and decreased the detected interaction for only a single sRNA (RybB).

**Figure 3.**
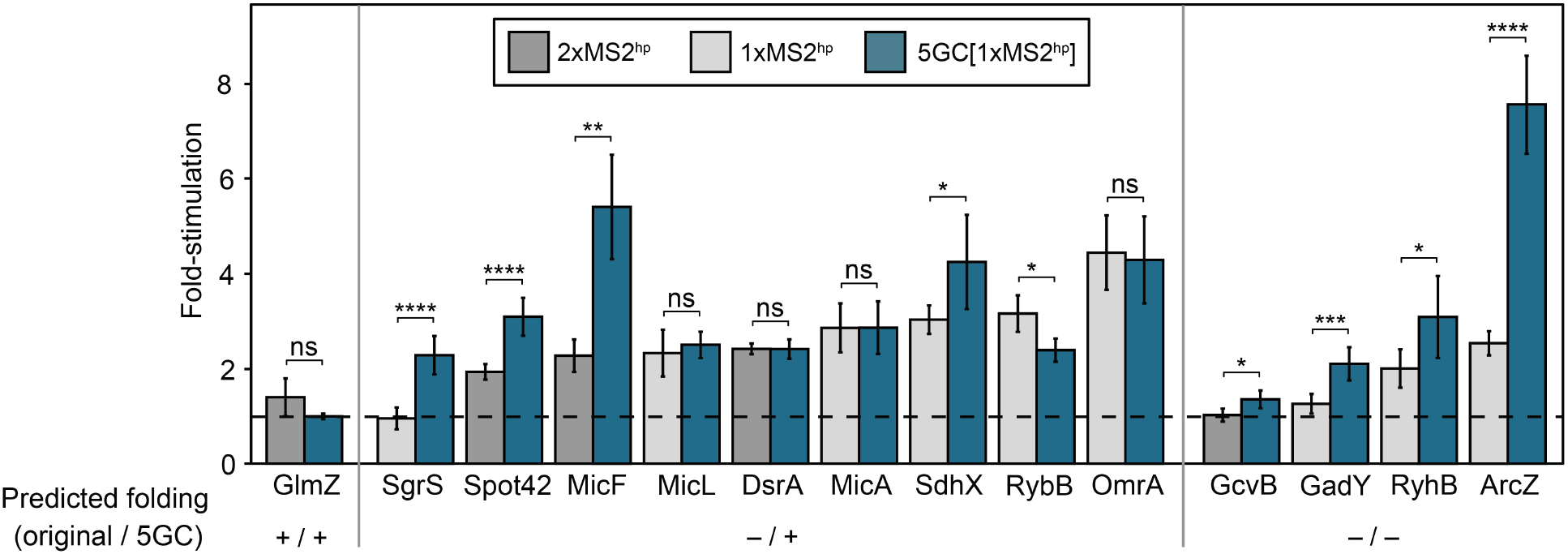
A short GC-clamp improves B3H signal for many Hfq-sRNA interactions. Results of β-gal assays performed in KB483 cells transformed with three compatible plasmids: pPrey that encoded α alone (–) or the α-Hfq fusion protein (pKB817), pAdapter that encoded λCI alone (–) or the CI-MS2^CP^ fusion protein (p35u4), and pBait that encoded a hybrid RNA with the indicated sRNA sequence following two copies (“2xMS2^hp^”; dark gray bars; pKB943, pKB858, pKB939, pKB941, or pKB940), one copy (“1xMS2^hp^”; light gray bars; pCH12, pLN97, pKB1212, pLN96, pKB1213, pLN94, pLN98, pCH10, or pCH13) of an MS2^hp^ moiety in the absence of a GC clamp, one copy of the MS2^hp^ moiety flanked by a 5bp GC clamp (“5GC”; pLN53, pLN91, pLN92, pLN85, pLN97, pLN54, pLN81, pLN96, pLN80, pLN94, pLN93, pLN98, or pLN84), or an RNA that contained only the MS2^hp^ moiety (pCH1 or pSS1). RNAs are grouped based on whether the hybrid RNAs are expected to fold correctly or incorrectly in the presence and absence of the 5GC clamp (see Supplemental Figs. S2-S6 for detailed sequences and predicted structures). Asterisks indicate P-values of student’s t-tests, represented as follows: ns, not significant, ^*^P < 0.05, ^**^P < 0.01, ^***^P < 0.001, ^****^P < 0.0001.

Secondary structure predictions suggest that the short GC clamp may differentially facilitate the proper folding of specific sRNAs, but no clear pattern emerges when comparing the effects of the GC clamp on B3H interaction to the predicted folding of pBait-sRNA constructs. For instance, of the nine pBait-sRNA constructs that were predicted to fold correctly *only* in the presence of the 5GC clamp (Supplemental Figs. S2, S4 and S5), the clamp improved Hfq B3H interactions for four RNAs (SgrS, Spot42, MicF, SdhX); had no significant effect on an additional four (MicA, MicL, DsrA, OmrA); and decreased the interaction for 1 RNA (RybB) (Fig. 3). On the other hand, four pBait-sRNA constructs were predicted to fold incorrectly in *either* the presence or absence of the GC clamp (GcvB, GadY, RyhB, ArcZ; Supplemental Figs. S3 and S6); nevertheless, the GC clamp improved B3H interactions for each of these RNAs (Fig. 3). Finally, the GlmZ pBait construct was predicted to fold correctly *regardless* of the presence of the GC clamp (Supplemental Fig. S3), but no B3H interaction was detectable with Hfq with either construct. The fact that the 172-nt GlmZ is longer than most of the other sRNAs used here may contribute to its undetectable B3H interaction with Hfq.

Overall, these results demonstrate the promise of a short GC clamp to expand detection of a broader set of sRNA targets in the B3H assay. The imperfect correlation between the predicted folding of these sRNAs and whether a B3H interaction with Hfq is detectable is consistent with the folding of a hybrid RNA not being the only limiting factor in detection of Hfq-sRNA interactions. Indeed, other considerations such as the stability of a particular hybrid transcript may also contribute to the detection of an RNA-protein interaction. In addition, computational predictions of secondary structure may not reflect the actual folding of the sRNAs *in vivo*, which can be determined by kinetics during transcription or interaction with cellular proteins, or can be constituted by an ensemble of variable conformations. Finally, for three of the eight RNAs that showed increased interactions with the 5GC clamp (Spot42, MicF, GcvB), this comparison was made to the original 2xMS2^hp^ constructs. Secondary structure predictions suggest that the 1xMS2^hp^ construct alone would have helped the folding of MicF, but not Spot42 or GcvB (Supplemental Figs. S2 and S3); however, we cannot distinguish between the contributions of the 5GC clamp and presence of a single MS2^hp^ in detecting interactions with these RNAs. Overall, since the 5GC[1xMS2^hp^] construct increases detectable interactions for many more RNAs than it hurts the interactions for, we recommend this pBait plasmid for cloning future RNA baits for detection in the B3H assay.

### Design of pBait constructs for RNA baits lacking an intrinsic terminator

All of the interactions above involve Hfq-dependent sRNAs that arise from the native 3′ end of their transcript and therefore encode their own intrinsic terminators. An important step to further expand the utility of the B3H assay is to develop constructs that can be used to express RNAs that do not provide their own terminator — *e*.*g*. 5′UTRs, ORF fragments, or heterologous RNAs from species with different transcription termination mechanisms. We selected *E. coli* Hfq-5′UTR interactions as a model system to develop and test constructs for B3H interactions with terminator-free RNAs. For expression of these 5′UTR-containing hybrid RNAs, we added an intrinsic terminator downstream of the HindIII restriction site in the 1xMS2^hp^ pBait construct (Fig. 4A). The terminator from the *E. coli trpA* transcript (T_*trpA*_) was selected due to its compact size of 28 nts. In addition to the terminator, we included a stop codon (UAA) so that any ribosomes that initiated translation on the RNA fragments containing a ribosomal binding site (RBS) and start codon in the hybrid RNA would also have a path to terminate translation. Initial secondary structure predictions suggested that certain RNAs like the *chiP* 5′UTR were likely to misfold in this 1xMS2^hp^-T_*trpA*_ construct (Fig. 4B). Given the success of GC clamps in supporting sRNA-Hfq B3H interactions, we explored their predicted effects on hybrid RNA folding in the context of T_*trpA*_-containing pBait constructs. In order to isolate bait RNAs from elements on both their 5′ and 3′ side (the MS2^hp^ and the intrinsic terminator, respectively), we designed constructs that placed the GC clamp on either side of the bait RNA itself (Fig. 4C). Secondary structure predictions suggested that the minimum length of GC clamp that promotes correct base pairing in these constructs was 7 bp (Fig. 4D), slightly longer than the 5 bp clamp that was sufficient for sRNA constructs above. While the 5′UTR of the *chiP* mRNA was predicted to fold incorrectly in the absence of a GC clamp, either a 7GC or 13GC clamp restored correct predicted folding (Fig. 4C,D). To test the effects of long and short GC clamps on the interaction of terminator-free RNAs, we selected 5′UTRs from *sodB, rpoS, eptB, chiP* and *mutS* mRNAs as well-characterized Hfq interactors (*e*.*g*., Mikulecky et al. 2004; Schu et al. 2015; Baba et al. 2006; Chen and Gottesman 2017; Geissmann and Touati 2004; Soper and Woodson 2008). Each 5′UTR fragment was cloned into 1xMS2^hp^-T*trpA* constructs or derivatives with a 7GC or 13GC clamp (Fig. 4A,C,D). Cloned fragments of these mRNAs began at transcription start sites and continued through the first 1-to-9 codons of the ORF (Supplemental Table S2), based on prior studies of mRNA-Hfq interactions (Mikulecky et al. 2004; Moon and Gottesman 2011; Schu et al. 2015).The predicted folding of all 5′UTR constructs is shown in Supplemental Figs. S7 and S8.

**Figure 4.**
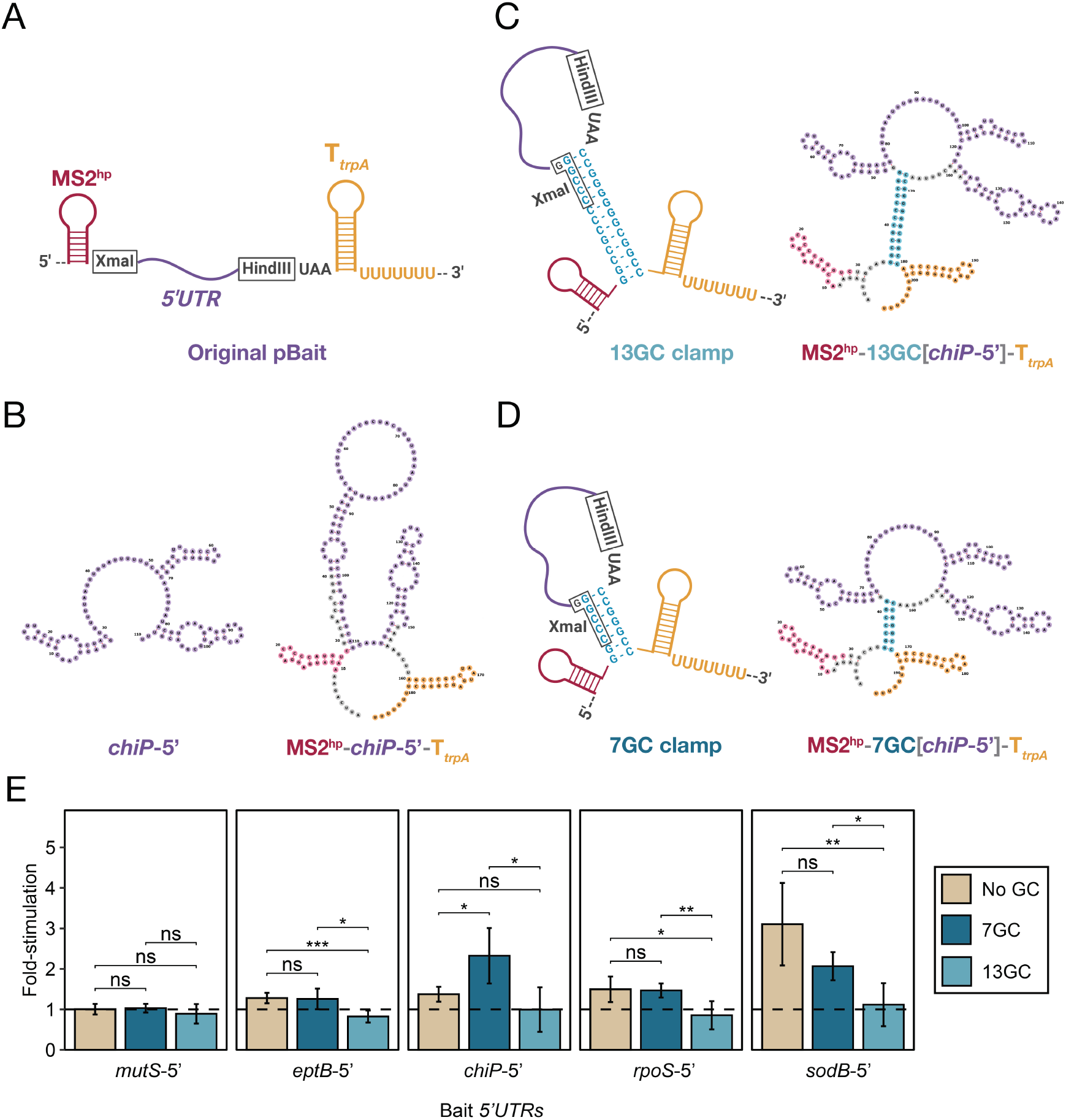
A short GC clamp improves B3H signal for RNAs that do not encode their own intrinsic terminator. (A) Schematic of a hybrid bait-RNA showing the intended secondary structure of MS2^hp^, a 5′UTR-containing RNA fragment inserted between the XmaI and HindIII restriction sites, and a *trpA* terminator (T_*trpA*_) (B) forna predictions of the *E. coli chiP* 5′UTR on its own (left) and when expressed as a hybrid RNA from the 1xMS2-T_*trpA*_ pBait plasmid (right). (C,D) Schematic of hybrid RNAs with additional GC clamps that are 13bp (C) or 5bp (D) long around the RNA bait. Schematics show the intended secondary structure (left) and forna-predicted secondary structure (right) of *chiP-5*′ expressed in these hybrid RNAs. (E) Results of β-gal assays performed in KB483 cells transformed with three compatible plasmids: pPrey that encoded α alone (–) or the α-Hfq fusion protein (pKB817), pAdapter that encoded λCI alone (–) or the CI-MS2^CP^ fusion protein (p35u4), and pBait that encoded a hybrid RNA with the 5′UTR of *E. coli* mRNAs *mutS, eptB, chiP, rpoS* or *sodB* inserted one copy of an MS2^hp^ moiety and the intrinsic terminator from *E. coli trpA* (T_trpA_) in the absence of a GC clamp (“No GC”; pHL40, pHL37, pHL39, pHL36, pHL38, respectively), or the presence of a GC clamp flanking the bait RNA moiety of either 7bp (“7GC”; pLN44, pLN42, pLN43, pLN41 or pSS3, respectively) or 13bp (“13GC”; pLN40, pLN38, pLN39, pLN37 or pSW3, respectively) or an RNA that contained only the MS2^hp^ moiety and *trpA* terminator (pHL6, pSS2 or pSW2). Student t-tests were conducted to assess statistical significance for differences between pairs of B3H interactions; asterisks indicate P-values, represented as follows: ns, not significant, *P < 0.05, ^**^P < 0.01, ^***^P< 0.001).

### A short GC clamp improves the detection of *chiP* 5′UTR-Hfq interaction

The interactions of the five selected 5′UTRs with Hfq were tested in the B3H system in the three versions of the T_*trpA*_-containing pBait construct. In the absence of a GC clamp (“no GC”), only *sodB*-5′ produced a reliably detectable signal in liquid β-gal assays (∼3-fold stimulation above background; Fig. 4D). In contrast, the signal for the interactions of *rpoS*-5′, *eptB*-5′, *chiP*-5′ and *mutS*-5′ RNAs with Hfq were less than 1.5-fold of basal *lacZ* activity (Fig. 4D). In the presence of a short GC clamp (“7GC”), there was a statistically significant increase in signal to ∼2-fold for *chiP-*5′ while the other interactions were not strongly affected (Fig. 4D). While the interaction of *sodB*-5′ with Hfq appears to decrease slightly in the presence of the short GC clamp, this effect is not statistically significant. Interestingly, however, the longer GC clamp results in undetectable signal (less than 1-fold) for most of the 5′UTR-Hfq interactions being tested. The observation that a short GC clamp enables the detection of two rather than one 5′UTR-Hfq interaction suggests that this 7GC clamp may indeed be helpful for B3H detection of RNA-protein interactions with RNAs that do not encode their own intrinsic terminator. However, as with the sRNA constructs, there are likely other limiting factors to the detection of 5′UTR-Hfq interactions, as three of the five interactions remained below 1.5-fold stimulation, even with the short GC clamp.

The intrinsic terminator within the 1xMS2^hp^-7GC[bait]-T_*trpA*_ pBait construct is likely helpful for creating a homogenous and stable hybrid RNA, and we recommend its use when performing B3H analysis on RNA baits that do not provide their own terminator, including non-bacterial RNAs. However, for any individual RNA-protein interaction, controls will be critical to ensure that the terminator itself does not contribute to interaction, especially when studying bacterial RBPs that have evolved to recognize intrinsic terminators. For instance, while the *trpA* terminator is not sufficient for binding to Hfq (Fig 4E), it may be that an interaction between the 5′UTR and the distal and/or rim surface of Hfq is further stabilized by a secondary interaction between the intrinsic terminator of *trpA* and Hfq’s proximal face. It will be interesting to explore this carefully in the future with mutagenesis studies, and to compare sequences and contexts of intrinsic terminators to minimize potential secondary interactions.

### Summary

In this work, we have developed several new pBait constructs for the B3H assay. The GC clamp and exogenous intrinsic terminator in these constructs are helpful to support the correct folding of hybrid RNAs and express RNAs that do not encode their own intrinsic terminators, respectively. We find that short GC clamps (5 or 7 bp long) are more effective than a longer 13bp GC clamp in the B3H assay. The new pBait constructs introduced here allow for detection of a broader range of RNA-protein interactions in B3H assay, with higher signal for many interactions. While there was a single sRNA-Hfq interaction that decreased in the presence of the 5GC clamp, overall, these short GC clamps increase detectable interactions for many more RNAs than they impair. We therefore recommend the use of constructs with a 5GC and/or 7GC-T_*trpA*_ as a starting point for exploring new bait RNAs in the B3H system.

## MATERIALS AND METHODS

### Bacterial Strains and Plasmid Construction

A complete list of plasmids, strains and oligonucleotides (oligos) used in this study is provided in Supplemental Tables S1–S3, respectively. NEB 5-α F′Iq cells (New England Biolabs) were used as the recipient strain for plasmid construction. KB483 (*O*_*L*_*2-62-lacZ; Δhfq*) (Pandey et al. 2020) was used as the reporter strain for all B3H experiments in this study. Strains and plasmids contain antibiotic-resistant genes as specified in Supplemental Tables S1 and S2, abbreviated as follows: TetR (tetracycline-resistant), KanR (kanamycin-resistant), StrR (streptomycin-resistant), CarbR (carbenicillin/ampicillin-resistant), CmR (chloramphenicol-resistant), and SpecR (spectinomycin-resistant).

Plasmids were constructed as specified in Supplemental Table S2 and the construction of key parent vectors is described below. Positive and negative control plasmids for new pBait constructs are available through Addgene, including pSS1 (pBait-5GC[1xMS2^hp^]), pSS2 (pBait-1xMS2^hp^-7GC-T_*trpA*_), pLN34 (pBait-5GC[1xMS2^hp^]-McaS), pLN43 (pBait-1xMS2^hp^-7GC [*chiP-*5′]-T_*trpA*_).

Plasmids pSW1 and pSS1 (pBait-13GC[1xMS2^hp^] and pBait-5GC[1xMS2^hp^], respectively) were constructed by PCR from partially complementary pairs of oligos (oLN28+29 and oLN30+31, respectively), which contained a GC clamp flanking the MS2^hp^ sequence along with EcoRI or XmaI restriction sites at either end. Following PCR and restriction enzyme digestion, these inserts were ligated into pCH1 (Pandey et al. 2020) digested with EcoRI and XmaI restriction enzymes. Plasmids pLN27 and pLN28 (pBait-13GC[1xMS2^hp^]-ChiX and pBait-5GC[1xMS2^hp^]-ChiX, respectively) were constructed analogously using the same pairs of oligos (oLN28+29 and oLN30+31, respectively), but inserted into pCH6 (Pandey et al. 2020).

The set of 5′UTR-containing hybrid RNA plasmids used in this study (pBait-1xMS2^hp^-5′UTR-TAA-T_*trpA*_, pHL36-40) were constructed from mutagenic PCR from their corresponding parent plasmids (pBait-1xMS2^hp^-5′UTR-T_*trpA*_; pHL28-32, respectively) to insert 3 nts encoding a stop codon (TAA) between the HindIII restriction site and the intrinsic terminator from the *E. coli trpA* transcript (T_*trpA*_).

Plasmids pSS2 and pSW2 (pBait-1xMS2^hp^-13GC-T_*trpA*_ and pBait-1xMS2^hp^-7GC-T_*trpA*_, respectively) were constructed from two rounds of mutagenic PCR from pHL6 (Pandey et al. 2020). The first round of site-directed PCR mutagenesis inserted the 5′ GC flank between the MS2^hp^ moiety and XmaI restriction site in pHL6, using end-to-end primers (oSW1+2 for 13GC insertion, resulting in pSW2x; oSS1+2 for 7GC insertion, resulting in pSS2x). These intermediate constructs were used as templates for the second round of mutagenesis to insert the 3′ GC flank along with 3 nts encoding a stop codon (TAA) between the HindIII restriction site and T_*trpA*_ sequence, using end-to-end primers (oSW3+4 for TAA-13GC insertion, resulting in pSW2; oSS3+4 for TAA-7GC insertion, resulting in pSS2). Plasmids pSW3 and pSS3 (pBait-1xMS2^hp^-13GC[*sodB-*5′]-T_*trpA*_ and pBait-1xMS2^hp^-7GC[*sodB-*5′]-T_*trpA*_, respectively) were constructed analogously. For the first round of mutagenesis, pHL38 (pBait-1xMS2^hp^-*sodB-* 5′TAA-T_*trpA*_) was used as a template plasmid for the insertion of the first GC flank using end-to-end primer pairs (oSW5+6 for 13GC insertion; oSS5+6 for 7 GC insertion) to create pSW3x and pSS3x, respectively. These intermediate constructs were then used as templates for the second round along with pairs of primers (oSW3+7 for 13GC insertion; oSS3+7 for 7GC insertion) to create the target plasmids (pSW3 and pSS3, respectively). PCR mutagenesis was conducted with Q5 Site-Directed Mutagenesis (New England Biolabs) using end-to-end primers designed with NEBaseChanger (Supplemental Table S3).

All other hybrid RNAs used in this study are constructed by inserting the RNA of interest into the XmaI/HindIII sites of the parent vectors for the original 2xMS2^hp^ / 1xMS2^hp^ pBait sRNA constructs (pKB845 / pCH1, respectively), the original 1xMS2^hp^ 5′UTR pBait construct (pHL6), and the long and short GC-clamp sRNA/5′UTR constructs (pSW1/pSS1 and pSW2/pSS2, respectively). More detailed information about all plasmids used in this study are provided in Supplementary Table S2.

### RNA structure predictions

Structure predictions of constructs used in this study were visualized by *forna*, an RNA secondary structure visualization tool using a force directed graph layout (http://rna.tbi.univie.ac.at/forna/) (Kerpedjiev et al. 2015). The predictions by forna were made based on the minimum free energy (MFE) structure calculated for the given sequence using RNAfold (http://rna.tbi.univie.ac.at/cgi-bin/RNAWebSuite/RNAfold.cgi) (Lorenz et al. 2011).

Note that figures throughout this work show sequences of hybrid RNAs that would arise from initiation at the purine nearest to the +1 position: the second A in the EcoRI site (GAATTC), but we have not empirically identified the transcription start site of the RNAs generated from pBait plasmids.

### B3H / β-galactosidase assays

For liquid B3H assays, reporter cells KB483 were transformed with three plasmids: pAdapter, expressing the CI-MS2^CP^ fusion protein (p35u4); pBait, expressing the hybrid RNA composed of the MS2^hp^ and the RNA of interest; and pPrey, expressing the protein of interest tethered to the N-terminal domain (NTD) of the α subunit of RNAP. Negative controls were transformed by replacing each of the hybrid components (pBait, pPrey and pAdapter), one at a time, with its “empty” version (MS2^hp^-empty for pBait, α-empty for pPrey, and CI-empty for pAdapter). Following heat shock to facilitate the uptake of plasmids, cells in each transformation were split in half (for duplicate setup) or thirds (for triplicate setup) and inoculated directly into liquid LB media supplemented with 0.2% arabinose (to induce RNA expression), 0-5 μM IPTG (to induce protein expression), and appropriate antibiotics to select for plasmids (Cm, Carb and Spec) and the reporter strain (Tet). Following overnight growth in a 96-deep-well plate at 37°C with shaking at 900 rpm, samples were back-diluted (1:40) into optically clear flat bottom 96-well plates containing LB media supplemented with the same concentration of antibiotics and arabinose as the overnight cultures, along with IPTG at 5, 10 or 15 μM. These back-dilutions were covered with sterile plastic lids and grown at 37°C to mid-log (OD_600_: 0.3-0.9; Molecular Devices SpectraMax) while shaking at 800-900 rpm. For each transformation, β-galactosidase (β-gal) activity was measured as previously described (Stockert et al. 2022), and was averaged across multiple replicates (duplicate or triplicate) carried out on the same day.

Three-hybrid interactions were quantified by calculating the fold-stimulation over basal β-gal levels for each interaction, defined as the β-gal value measured in the presence of all hybrid constructs divided by the highest activity from negative control transformations in which part of each hybrid component is missing (-bait, -adapter or -prey). Bar plots represent the mean and standard deviations of fold-stimulation (SD) values collected across at least three independent experiments. Statistical analyses were performed using two-tailed, unpaired, Student’s t-tests. P values are represented with the follow abbreviations: ns, not significant, ^****^P < 0.0001, ^***^P < 0.001, ^**^P < 0.01 and ^*^P < 0.05.

## ACKNOWLEDGEMENTSS

This work was supported by the National Institutes of Health [2R15GM135878], the Henry R. Luce foundation, the Camille and Henry Dreyfus Foundation, and Mount Holyoke College. We thank members of the Berry Lab and Amy Camp for helpful discussions and feedback; Suxuan Wang and Sylvie Schein for help with construct design; and Courtney Hegner and Chandra Gravel for assistance with cloning.

## AUTHOR CONTRIBUTIONS

L.D.N and H.L. designed and cloned plasmids and tested them in B3H assays. K.E.B. conceived of the project, obtained funding, and oversaw the experimental implementation. L.D.N, H.L. and K.E.B analyzed data and L.D.N. completed statistical analyses. L.D.N. and K.E.B made the figures and wrote the manuscript with input and feedback from H.L.

## SUPPLEMENTAL MATERIAL

Supplemental materials for this article are available online, including Supplemental Tables S1 to S3, and Supplemental Figures S1 to S8.

