## Supplemental Information for "Improved constructs for bait RNA display in a bacterial three-hybrid assay"

This pdf file includes:

Supplemental Tables S1 to S3

Supplemental Figures S1 to S8

Supplemental References

**Supplemental Table S1.** *E. coli* strains used in this study.

| **Strain** | **Genotype** | **Antibiotic Resistance** | **Source** |
|---|---|---|---|
| NEB5α-F′$I^q$ | *E. coli* $lacI^q$ host strain for plasmid construction. | TetR | New England Biolabs |
| KB483 | FW102 Δ*hfq::kan* harboring an F′ episome bearing test promoter *plac-O$_L$2-62* fused to *lacZ*. | KanR; TetR; StrR | (Pandey et al. 2020) |
| MG1655 | Strain K-12 F⁻ lambda⁻ *ivlG*⁻ *rfb*-50 *rph*-1 | – | (Blattner et al. 1997) |

**Supplemental Table S2.** Plasmids used in this study.

| **Plasmid Name** | **Description** | **Details** | **Source or oligos used in this study** |
|---|---|---|---|
| p35u4 (Addgene #174786) | pAdapter: pAC-$p_{constit}$-CI-$MS2^{CP}$ | Encodes CI-$MS2^{CP}$ fusion protein under control of a constitutive promoter; p15A origin of replication. Isolated from a pCW17-derived mutagenesis library; confers CmR | (Wang et al. 2021) |
| pACλCI (Addgene #53730) | pACλCI empty vector | Encodes full-length λCI under the control of the lac*UV5* promoter; confers CmR | (Pandey et al. 2020) |
| pBr-α (Addgene #53731) | pBR-α-empty vector | Encodes residues 1-248 of the alpha vector fused under control of *lpp* and *lacUV5 promoters*; confers AmpR | (Dove et al. 1997) |
| pCH1 (Addgene #174663) | pBait-1x$MS2^{hp}$-empty | pCDF-pBAD-1x$MS2^{hp}$. Encodes a single $MS2^{hp}$ under the control of an arabinose-inducible promoter, followed by XmaI and HindIII sites. CloDF13 origin of replication; confers SpecR | (Pandey et al. 2020) (schematics in Fig 1B+2A) |
| pCH6 (Addgene #174662) | pBait-1x$MS2^{hp}$-ChiX | *E. coli chiX* inserted between XmaI/HindIII sites in pCH1; sRNA encodes its own terminator; confers SpecR | (Pandey et al. 2020) |
| pCH7 | pBait-1x$MS2^{hp}$-McaS | *E. coli mcaS* inserted between XmaI/HindIII sites in pCH1; sRNA encodes its own terminator; confers SpecR | oKB1192+ oKB1193 |
| pCH8 | pBait-1x$MS2^{hp}$-CyaR | *E. coli cyaR* inserted between XmaI/HindIII sites in pCH1; sRNA encodes its own terminator; confers SpecR | oKB1194+ oKB1195 |
| pCH9 | pBait-1x$MS2^{hp}$-OxyS | *E. coli oxyS* inserted between XmaI/HindIII sites in pCH1; sRNA encodes its own terminator; confers SpecR | (Pandey et al. 2020) |

| | | | |
|---|---|---|---|
| pCH10 | pBait-1xMS2$^{hp}$-RyhB | *E. coli ryhB* inserted between XmaI/HindIII sites in pCH1; sRNA encodes its own terminator; confers SpecR | oKB1198+ oKB1199 |
| pCH12 | pBait-1xMS2$^{hp}$-SgrS | *E. coli sgrS* inserted between XmaI/HindIII sites in pCH1; sRNA encodes its own terminator; confers SpecR | oKB1107+ oKB1108 |
| pCH13 | pBait-1xMS2$^{hp}$-ArcZ | *E. coli arcZ* inserted between XmaI/HindIII sites in pCH1; sRNA encodes its own terminator; confers SpecR | (Pandey et al. 2020) |
| pHL6 | pBait-1xMS2$^{hp}$-T$_{trpA}$ | Encodes a single MS2$^{hp}$ followed by XmaI and HindIII sites and the intrinsic terminator from *E. coli trpA* gene (T$_{trpA}$). Hybrid RNA is under the control of an arabinose-inducible promoter. CloDF13 origin of replication; confers SpecR | (Pandey et al. 2020) (schematic: Fig 4A) |
| pHL28 | pBait-1xMS2$^{hp}$-*rpoS*-5′-T$_{trpA}$ | *E. coli* 5′UTR of *rpoS* (-134 to +3, relative to AUG) inserted between XmaI/HindIII sites in pHL6; confers SpecR | oHL38 + oHL39 |
| pHL29 | pBait-1xMS2$^{hp}$-*eptB*-5′-T$_{trpA}$ | *E. coli* 5′UTR of *eptB* (-106 to +27, relative to AUG) inserted between XmaI/HindIII sites in pHL6; confers SpecR | oHL40 + oHL41 |
| pHL30 | pBait-1xMS2$^{hp}$-*sodB*-5′-T$_{trpA}$ | *E. coli* 5′UTR of *sodB* (-55 to +27, relative to AUG) inserted between XmaI/HindIII sites in pHL6; confers SpecR | oHL42 + oHL43 |
| pHL31 | pBait-1xMS2$^{hp}$-*chiP*-5′-T$_{trpA}$ | *E. coli* 5′UTR of *chiP* (-98 to +12, relative to AUG) inserted between XmaI/HindIII sites in pHL6; confers SpecR | oHL44 + oHL45 |
| pHL32 | pBait-1xMS2$^{hp}$-*mutS*-5′-T$_{trpA}$ | *E. coli* 5′UTR of *mutS* (-74 to +27, relative to AUG) inserted between XmaI/HindIII sites in pHL6; confers SpecR | oHL46 + oHL47 |
| pHL36 | pBait-1xMS2$^{hp}$-*rpoS*-5′-TAA-T$_{trpA}$ | pHL28 with stop codon (TAA) inserted after 5′UTR: *E. coli* 5′UTR of *mutS* (-134 to +3, relative to AUG), upstream of T$_{trpA}$; confers SpecR | oHL66 + oHL67 |
| pHL37 | pBait-1xMS2$^{hp}$-*eptB*-5′-TAA-T$_{trpA}$ | pHL29 with stop codon (TAA) inserted after 5′UTR: *E. coli* 5′UTR of *eptB* (-106 to +27, relative to AUG), upstream of T$_{trpA}$; confers SpecR | oHL66 + oHL68 |
| pHL38 | pBait-1xMS2$^{hp}$-*sodB*-5′TAA-T$_{trpA}$ | pHL30 with stop codon (TAA) inserted after 5′UTR: *E. coli* 5′UTR of *sodB* (-55 to +27, relative to AUG), upstream of T$_{trpA}$; confers SpecR | oHL66 + oHL69 |

| | | | |
|---|---|---|---|
| pHL39 | pBait-1xMS2$^{hp}$-*chiP*-5′-TAA-$T_{trpA}$ | pHL31 with stop codon (TAA) inserted between 5′UTR and HindIII site: *E. coli* 5′UTR of *chiP* (-98 to +12, relative to AUG) upstream of $T_{trpA}$; confers SpecR | oHL66 + oHL70 |
| pHL40 | pBait-1xMS2$^{hp}$-*mutS*-5′-TAA-$T_{trpA}$ | pHL32 with stop codon (TAA) inserted between 5′UTR and HindIII site: *E. coli* 5′UTR of *mutS* (-74 to +27, relative to AUG) upstream of $T_{trpA}$; confers SpecR | oHL66 + oHL71 |
| pKB817 | pPrey-α-Hfq | Encodes residues 1-248 of the alpha subunit of RNA polymerase fused via three alanine residues to full-length wild-type *E. coli hfq*; confers AmpR | (Berry and Hochschild 2018) |
| pKB845 | pBait-2xMS2$^{hp}$ | pCDF-pBAD-2xMS2$^{hp}$-XmaI-HindIII; two MS2 RNA hairpins (2xMS2$^{hp}$) and an XmaI site inserted into pKB822 between BamHI/HindIII sites; confers SpecR | (Berry and Hochschild 2018) |
| pKB858 | pBait-2xMS2$^{hp}$-Spot42 | *E. coli spot42* inserted between XmaI/HindIII sites in pKB845; sRNA encodes its own terminator; confers SpecR | oKB1113 + oKB1114 |
| pKB939 | pBait-2xMS2$^{hp}$-MicF | *E. coli micF* inserted between XmaI/HindIII sites in pKB845; sRNA encodes its own terminator; confers SpecR | oKB1205 + oKB1206 |
| pKB940 | pBait-2xMS2$^{hp}$-GcvB | *E. coli gcvB* inserted between XmaI/HindIII sites in pKB845; sRNA encodes its own terminator; confers SpecR | oKB1207 + oKB1208 |
| pKB941 | pBait-2xMS2$^{hp}$-DsrA | *E. coli dsrA* inserted between XmaI/HindIII sites in pKB845; sRNA encodes its own terminator; confers SpecR | (Wang et al. 2021) |
| pKB943 | pBait-2xMS2$^{hp}$-GlmZ | *E. coli glmZ* inserted between XmaI/HindIII sites in pKB845; sRNA encodes its own terminator; confers SpecR | oKB1213 + oKB1214 |
| pKB1212 | pBait-1xMS2$^{hp}$-MicA | *E. coli micA* inserted between XmaI/HindIII sites in pKB845; sRNA encodes its own terminator; confers SpecR | oKB1535 + oKB1536 |
| pKB1213 | pBait-1xMS2$^{hp}$-RybB | *E. coli rybB* inserted between XmaI/HindIII sites in pKB845; sRNA encodes its own terminator; confers SpecR | oKB1537 + oKB1538 |
| pLN27 | pBait-13GC[1xMS2$^{hp}$]-ChiX | pCH6 with 13-bp GC clamp flanking an MS2$^{hp}$ inserted between EcoRI/XmaI sites; sRNA encodes its own terminator; confers SpecR | pCH6 + oLN28 + oLN29 |
| pLN28 | pBait-5GC[1xMS2$^{hp}$]-ChiX | pCH6 with 5-bp GC clamp flanking an MS2$^{hp}$ inserted between EcoRI/XmaI sites; sRNA encodes its own terminator; confers SpecR | pCH6 + oLN30 + oLN31 |

| | | | |
|---|---|---|---|
| pLN31 | pBait-13GC[1xMS2$^{hp}$]-McaS | *E. coli mcaS* inserted between XmaI/HindIII sites in pSW1; sRNA encodes its own terminator; confers SpecR | oKB1192 + oKB1193 |
| pLN32 | pBait-13GC[1xMS2$^{hp}$]-CyaR | *E. coli cyaR* inserted between XmaI/HindIII sites in pSW1; sRNA encodes its own terminator; confers SpecR | oKB1194 + oKB1195 |
| pLN33 | pBait-13GC[1xMS2$^{hp}$]-OxyS | *E. coli oxyS* inserted between XmaI/HindIII sites in pSW1; sRNA encodes its own terminator; confers SpecR | oKB1196 + oKB1197 |
| pLN34 (Addgene #222403) | pBait-5GC[1xMS2$^{hp}$]-McaS | *E. coli mcaS* inserted between XmaI/HindIII sites in pSS1; sRNA encodes its own terminator; confers SpecR | oKB1192 + oKB1193 |
| pLN35 | pBait-5GC[1xMS2$^{hp}$]-CyaR | *E. coli cyaR* inserted between XmaI/HindIII sites in pSS1; sRNA encodes its own terminator; confers SpecR | oKB1194 + oKB1195 |
| pLN36 | pBait-5GC[1xMS2$^{hp}$]-OxyS | *E. coli oxyS* inserted between XmaI/HindIII sites in pSS1; sRNA encodes its own terminator; confers SpecR | oKB1196 + oKB1197 |
| pLN37 | pBait-1xMS2$^{hp}$-13GC[*rpoS*-5′]-$T_{trpA}$ | *E. coli* 5′UTR of *rpoS* (-134 to +3, relative to AUG; Mikulecky et al. 2004) inserted between XmaI/HindIII sites in pSW2; confers SpecR | oHL38 + oHL39 |
| pLN38 | pBait-1xMS2$^{hp}$-13GC[*eptB*-5′] -$T_{trpA}$ | *E. coli* 5′UTR of *eptB* (-106 to +27, relative to AUG) inserted between XmaI/HindIII sites in pSW2; confers SpecR | oHL40 + oHL41 |
| pLN39 | pBait-1xMS2$^{hp}$-13GC[*chiP*-5′]-$T_{trpA}$ | *E. coli* 5′UTR of *chiP* (-98 to +12, relative to AUG) inserted between XmaI/HindIII sites in pSW2; confers SpecR | oHL44 + oHL45 |
| pLN40 | pBait-1xMS2$^{hp}$-13GC[*mutS*-5′]-$T_{trpA}$ | *E. coli* 5′UTR of *mutS* (-74 to +27, relative to AUG) inserted between XmaI/HindIII sites in pSW2; confers SpecR | oHL46 + oHL47 |
| pLN41 | pBait-1xMS2$^{hp}$-7GC[*rpoS*-5′]-$T_{trpA}$ | *E. coli* 5′UTR of *rpoS* (-134 to +3, relative to AUG; Mikulecky et al. 2004) inserted between XmaI/HindIII sites in pSS2; confers SpecR | oHL38 + oHL39 |
| pLN42 | pBait-1xMS2$^{hp}$-7GC[*eptB*-5′]-$T_{trpA}$ | *E. coli* 5′UTR of *eptB* (-106 to +27, relative to AUG) inserted between XmaI/HindIII sites in pSS2; confers SpecR | oHL40 + oHL41 |
| pLN43 (Addgene #222404) | pBait-1xMS2$^{hp}$-7GC[*chiP*-5′]-$T_{trpA}$ | *E. coli* 5′UTR of *chiP* (-98 to +12, relative to AUG) inserted between XmaI/HindIII sites in pSS2; confers SpecR | oHL44 + oHL45 |
| pLN44 | pBait-1xMS2$^{hp}$-7GC[*mutS*-5′]-$T_{trpA}$ | *E. coli* 5′UTR of *mutS* (-74 to +27, relative to AUG) inserted between XmaI/HindIII sites in pSS2; confers SpecR | oHL46 + oHL47 |

| | | | |
|---|---|---|---|
| pLN53 | pBait-5GC[1xMS2$^{hp}$]-GlmZ | *E. coli glmZ* inserted between XmaI/HindIII sites in pSS1; sRNA encodes its own terminator; confers SpecR | oKB1213 + oKB1214 |
| pLN54 | pBait-5GC[1xMS2$^{hp}$]-DsrA | *E. coli dsrA* inserted between XmaI/HindIII sites in pSS1; sRNA encodes its own terminator; confers SpecR | oKB1209 + oKB1210 |
| pLN80 | pBait-5GC[1xMS2$^{hp}$]-RybB | *E. coli rybB* inserted between XmaI/HindIII sites in pSS1; sRNA encodes its own terminator; confers SpecR | oKB1537 + oKB1538 |
| pLN81 | pBait-5GC[1xMS2$^{hp}$]-MicA | *E. coli micA* inserted between XmaI/HindIII sites in pSS1; sRNA encodes its own terminator; confers SpecR | oKB1535 + oKB1536 |
| pLN83 | pBait-5GC[1xMS2$^{hp}$]-RyhB | *E. coli ryhB* inserted between XmaI/HindIII sites in pSS1; sRNA encodes its own terminator; confers SpecR | oKB1198 + oKB1199 |
| pLN84 | pBait-5GC[1xMS2$^{hp}$]-ArcZ | *E. coli arcZ* inserted between XmaI/HindIII sites in pSS1; sRNA encodes its own terminator; confers SpecR | oKB1211 + oKB1212 |
| pLN85 | pBait-5GC[1xMS2$^{hp}$]-MicF | *E. coli micF* inserted between XmaI/HindIII sites in pSS1; sRNA encodes its own terminator; confers SpecR | oKB1205 + oKB1206 |
| pLN91 | pBait-5GC[1xMS2$^{hp}$]-SgrS | *E. coli sgrS* inserted between XmaI/HindIII sites in pSS1; sRNA encodes its own terminator; confers SpecR | oKB1107 + oKB1108 |
| pLN92 | pBait-5GC[1xMS2$^{hp}$]-Spot42 | *E. coli spot42* inserted between XmaI/HindIII sites in pSS1; native sRNA terminator; confers SpecR | oKB1113 + oKB1114 |
| pLN93 | pBait-5GC[1xMS2$^{hp}$]-GcvB | *E. coli gcvB* inserted between XmaI/HindIII sites in pSS1; sRNA encodes its own terminator; confers SpecR | oKB1207 + oKB1208 |
| pLN94 | pBait-1xMS2$^{hp}$-OmrA | *E. coli omrA* inserted between XmaI/HindIII sites in pSS1; sRNA encodes its own terminator; confers SpecR | oLN66 + oLN67 |
| pLN96 | pBait-1xMS2$^{hp}$-SdhX | *E. coli sdhX* inserted between XmaI/HindIII sites in pSS1; sRNA encodes its own terminator; confers SpecR | oLN70 + oLN71 |
| pLN97 | pBait-1xMS2$^{hp}$-MicL | *E. coli micL* inserted between XmaI/HindIII sites in pSS1; sRNA encodes its own terminator; confers SpecR | oLN72 + oLN73 |
| pLN98 | pBait-1xMS2$^{hp}$-GadY | *E. coli gadY* inserted between XmaI/HindIII sites in pSS1; sRNA encodes its own terminator; confers SpecR | oLN74 + oLN75 |

| | | | |
|---|---|---|---|
| pLN100 | pBait-5GC[1xMS2$^{hp}$]-OmrA | *E. coli omrA* inserted between XmaI/HindIII sites in pSS1; sRNA encodes its own terminator; confers SpecR | oLN66 + oLN67 |
| pLN102 | pBait-5GC[1xMS2$^{hp}$]-SdhX | *E. coli sdhX* inserted between XmaI/HindIII sites in pSS1; sRNA encodes its own terminator; confers SpecR | oLN70 + oLN71 |
| pLN103 | pBait-5GC[1xMS2$^{hp}$]-MicL | *E. coli micL* inserted between XmaI/HindIII sites in pSS1; sRNA encodes its own terminator; confers SpecR | oLN72 + oLN73 |
| pLN104 | pBait-5GC[1xMS2$^{hp}$]-GadY | *E. coli gadY* inserted between XmaI/HindIII sites in pSS1; sRNA encodes its own terminator; confers SpecR | oLN74 + oLN75 |
| pSS1 (Addgene #222405) | pBait-5GC[1xMS2$^{hp}$] | pCDF-pBAD-5GC[1xMS2$^{hp}$]. Encodes a single MS2$^{hp}$ flanked by 5-bp GC clamp, followed by XmaI and HindIII sites; hybrid RNA is under the control of an arabinose-inducible promoter, CloDF13 origin of replication; confers SpecR | pCH1 + oLN30 + oLN31 (schematic in Fig 2D) |
| pSS2 (Addgene #222406) | pBait-1xMS2$^{hp}$-7GC -$T_{trpA}$ | pCDF-pBAD-1xMS2$^{hp}$-7GC[XmaI-HindIII]-TAA-$T_{trpA.}$ Encodes a single MS2$^{hp}$ followed by XmaI and HindIII sites flanked by a 7-bp GC clamp, a stop codon (TAA) and the intrinsic terminator from *E. coli trpA* gene ($T_{trpA}$). Hybrid RNA is under the control of an arabinose-inducible promoter. CloDF13 origin of replication; confers SpecR | pSS2x + oSS3 + oSS4 (schematic in Fig 4D) |
| pSS2x | n/a | Intermediate construct used to make pSS2 from pHL6 | pHL6 + oSS1 + oSS2 |
| pSS3 | pBait-1xMS2$^{hp}$-7GC[*sodB*-5′]-$T_{trpA}$ | *E. coli* 5′UTR of *sodB* (-55 to +27, relative to AUG) between XmaI/HindIII sites in pSS2 (cloned by inserting 7GC clamp into pHL38); confers SpecR | pSS3x + oSS3 + oSS7 |
| pSS3x | n/a | Intermediate construct used to make pSS23 from pHL38 | pHL30 + oSS5 + oSS6 |
| pSW1 | pBait-1xMS2$^{hp}$-13GC[1xMS2$^{hp}$] | pCDF-pBAD-13GC[1xMS2$^{hp}$]. Encodes a single MS2$^{hp}$ flanked by 13-bp GC clamp, followed by XmaI and HindIII sites; hybrid RNA is under the control of an arabinose-inducible promoter, CloDF13 origin of replication; confers SpecR | oLN28 + oLN29 (schematic in Fig 2C) |
| pSW2 | pBait-1xMS2$^{hp}$-13GC-$T_{trpA}$ | pCDF-pBAD-1xMS2$^{hp}$-13GC[XmaI-HindIII]-TAA-$T_{trpA}$ Encodes a single MS2$^{hp}$ followed by XmaI and HindIII sites flanked by a 13-bp GC clamp, a stop codon (TAA) and the | pSW2x + oSW3 + oSW4 (schematic in Fig 4C) |

| | | | |
|---|---|---|---|
| | | intrinsic terminator from *E. coli trpA* gene ($T_{trpA}$). Hybrid RNA is under the control of an arabinose-inducible promoter. CloDF13 origin of replication; confers SpecR | |
| pSW2x | n/a | Intermediate construct used to make pSW2 from pHL6 | pHL6 + oSW1+ oSW2 |
| pSW3 | pBait-1xMS2$^{hp}$-13GC[*sodB*-5′]-$T_{trpA}$ | pHL38 with 13-bp GC clamp flanking *E. coli* 5′UTR of *sodB* and stop codon (TAA) inserted between HindIII site and the second GC flank; confers SpecR | pSW3x + oSW3 + oSW7 |
| pSW3x | n/a | Intermediate construct used to make pSW3 from pHL38 | pHL30 + oSW5 + oSW6 |

**Supplemental Table S3.** Oligonucleotides used in this study.

| **Oligo Name** | **Sequence (5′ to 3′)** | **Details (F: Forward primer, R: Reverse primer)** |
|---|---|---|
| oHL38 | GGCCGGCCCGGGACACGCTTGCATTTTGAAATTCG | F XmaI *rpoS*-5′ (pHL28, pLN37, pLN41) |
| oHL39 | GGCCGGAAGCTTCATAAGGTGGCTCCTACCCGTGATC | R HindIII *rpoS*-5′ (pHL28, pLN37, pLN41) |
| oHL40 | GGCCGGCCCGGGGCGCGTGTAGATTTTACTTATCTGAC | F XmaI *eptB*-5′ (pHL29, pLN38, pLN42) |
| oHL41 | GGCCGGAAGCTTCTGTGTAATCGATTTGATGTATCTCATG | R HindIII *eptB*-5′ (pHL29, pLN38, pLN42) |
| oHL42 | GGCCGGCCCGGGATACGCACAATAAGGCTATTGTACG | F XmaI *sodB*-5′ (pHL30) |
| oHL43 | GGCCGGAAGCTTTGGTAGTGCAGGTAATTCGAATGAC | R HindIII *sodB*-5′ (pHL30) |
| oHL44 | GGCCGGCCCGGGGTAGTCAGCGAGACTTTTCTCAACGC | F XmaI *chiP*-5′ (pHL31, pLN39, pLN43) |
| oHL45 | GGCCGGAAGCTTAAACGTACGCATGGGTTAATCCTCTTTG | R HindIII *chiP*-5′ (pHL31, pLN39, pLN43) |
| oHL46 | GGCCGGCCCGGGTGCGCCTTATGTGATTACAACGAAA ATA | F XmaI *mutS*-5′ (pHL32, pLN40, pLN44) |
| oHL47 | GGCCGGAAGCTTGGCGTCGAAATTTTCTATTGCACTC | R HindIII *mutS*-5′ (pHL32, pLN40, pLN44) |
| oHL66 | TAAAAGCTTAGCCCGCCTAAT | F Q5 TAA (pHL36, 37, 38, 39, 40) |

| | | |
|---|---|---|
| oHL67 | CATAAGGTGGCTCCTACC | R Q5 *rpoS*-5′ TAA (pHL36) |
| oHL68 | CTGTGTAATCGATTTGATGTATCTC | R Q5 *eptB*-5′ TAA (pHL37) |
| oHL69 | TGGTAGTGCAGGTAATTCG | R Q5 *sodB*-5′ TAA (pHL38) |
| oHL70 | AAACGTACGCATGGGTTAATC | R Q5 *chiP*-5′ TAA (pHL39) |
| oHL71 | GGCGTCGAAATTTTCTATTGC | R Q5 *mutS*-5′ TAA (pHL40) |
| oKB1107 | TCCCCCCGGGGATGAAGCAAGGGGGTGCC | F XmaI SgrS (pCH12, pLN91) |
| oKB1108 | CCGGCCAAGCTTAAAAAAAACCAGCAGGTATAATCTGCTGG | R HindIII SgrS (pCH12, pLN91) |
| oKB1192 | TCCCCCCGGGACCGGCGCAGAGGAG | F XmaI McaS (pCH7, pLN31, pLN34) |
| oKB1193 | CCGGCCAAGCTTAAAAAATAGAGTCTGTCGACATCCGCC | R HindIII McaS (pCH7, pLN31, pLN34) |
| oKB1194 | TCCCCCCGGGGCTGAAAAACATAACCCATAAAATGCTAGC | F XmaI CyaR (pCH8, pLN32, pLN35) |
| oKB1195 | CCGGCCAAGCTTAAAAAATAAGCCCGTGTAAGGGAGATTAC | R HindIII CyaR (pCH8, pLN32, pLN35) |
| oKB1196 | TCCCCCCGGGGAAACGGAGCGGCACCTC | F XmaI OxyS (pCH9, pLN33, pLN36) |
| oKB1197 | CCGGCCAAGCTTAAAAAAAAGCGGATCCTGGAGATCC | R HindIII OxyS (pCH9, pLN33, pLN36) |
| oKB1198 | TCCCCCCGGGGCGATCAGGAAGACCCTCG | F XmaI RyhB (pCH10, pLN83) |
| oKB1199 | CCGGCCAAGCTTAAAAAAAAGCCAGCACCCGG | R HindIII RyhB (pCH10, pLN83) |
| oKB1205 | TCCCCCCGGGGCTATCATCATTAACTTTATTTATTACCGTC | F XmaI MicF (pKB939, pLN85) |
| oKB1206 | CCGGCCAAGCTTAAAAAAAACCGAATGCGAGGCATC | R HindIII MicF (pKB939, pLN85) |
| oKB1207 | TCCCCCCGGGACTTCCTGAGCCGGAACG | F XmaI GcvB (pKB940, pLN93) |
| oKB1208 | CCGGCCAAGCTTAAAAAAAGCACCGCAATTAGGCG | R HindIII GcvB (pKB940, pLN93) |

| | | |
|---|---|---|
| oKB1209 | TCCCCCCGGGAACACATCAGATTTCCTGGTGTAAC | F XmaI DsrA (pKB941, pLN54) |
| oKB1210 | CCGGCCAAGCTTAAAAAAAATCCCGACCCTGAGGG | R HindIII DsrA (pKB941, pLN54) |
| oKB1211 | TCCCCCCGGGGTGCGGCCTGAAAAACAGTGC | F XmaI ArcZ (pCH13, pLN84) |
| oKB1212 | CCGGCCAAGCTTAAAAAATGACCCCGGCTAGACC | R HindIII ArcZ (pCH13, pLN84) |
| oKB1213 | TCCCCCCGGGGTAGATGCTCATTCCATCTCTTATGTTCG | F XmaI GlmZ (pKB943, pLN53) |
| oKB1214 | CCGGCCAAGCTTAAAAAAAACAGGTCTGTATGACAACAAGTGG | R HindIII GlmZ (pKB943, pLN53) |
| oKB1535 | GGCCGGCCCGGGGAAAGACGCGCATTTGTTATCATCATCC | F XmaI MicA (pKB1212, pLN81) |
| oKB1536 | CCGGCCAAGCTTAGAAAAGAAAAAGGCCACTCGTGAGTG | R HindIII MicA (pKB1212, pLN81) |
| oKB1537 | GGCCGGCCCGGGGCCACTGCTTTTCTTTGATGTCCCC | F XmaI RybB (pKB1213, pLN80) |
| oKB1538 | CCGGCCAAGCTTAACAAAAAACCCATCAACCTTGAACCG | R HindIII RybB (pKB1213, pLN80) |
| oLN28 | AATTCACCGGGCGCGGCGCAGAAAACATGAGGATCACCCATGTCTGCAGGCGCCGCGC | F EcoRI 13GC (pLN27, pSW1) |
| oLN29 | CCGGGCGCGGCGCCTGCAGACATGGGTGATCCTCATGTTTTCTGCGCCGCGCCCGGTG | R XmaI 13GC (pLN27, pSW1) |
| oLN30 | AATTCACCGGGAGAAAACATGAGGATCACCCATGTCTGCAGC | F EcoRI 5GC (pLN28, pSS1) |
| oLN31 | CCGGGCTGCAGACATGGGTGATCCTCATGTTTTCTCCCGGTG | R XmaI 5GC (pLN28, pSS1) |
| oLN66 | AAGCTTACAGAATTTTAAGTGCTTC | F XmaI OmrA (pLN94, pLN100) |
| oLN67 | ATTAGGTGACATCACGAAGG | R HindIII OmrA (pLN94, pLN100) |
| oLN70 | TCCCCCCGGGATATCTGTAATAAGAAATAGCCCTCGCC | F XmaI SdhX (pLN96, pLN102) |
| oLN71 | CCGGCCAAGCTTACAAAAAAGGCCATCATACGATGG | R HindIII SdhX (pLN96, pLN102) |

| | | |
|---|---|---|
| oLN72 | TCCCCCCGGGATTTTTACCGTTGCATCATGTCGC | F XmaI MicL (pLN97, pLN103) |
| oLN73 | CCGGCCAAGCTTAAAAAAAGGCCCCTGTTGAAATTGC | R HindIII MicL (pLN97, pLN103) |
| oLN74 | TCCCCCCGGGACTGAGAGCACAAAGTTTCCCG | F XmaI GadY (pLN98, pLN104) |
| oLN75 | CCGGCCAAGCTTAAAAAAAACCCGGCATAGGGGAC | R HindIII GadY (pLN98, pLN104) |
| oSS1 | GGCCCGGGACCTGCAGGCAT | F Q5 7GC (flank 1) (pSS2x) |
| oSS2 | CTGCAGACATGGGTGATCCTCATG | R Q5 7GC (flank 1) (pSS2x) |
| oSS3 | GGGCCAGCCCGCCTAATGAGCGG | F Q5 7GC (flank 2)-TAA (pSS2, pSS3) |
| oSS4 | GGTTAAAGCTTGCATGCCTGCAGG | R Q5 7GC (flank 2)-TAA (pSS2) |
| oSS5 | GGCCCGGGATACGCACAATA | F Q5 7GC (flank 1) (pSS3x) |
| oSS6 | CTGCAGACATGGGTGATC | R Q5 7GC (flank 1) (pSS3x) |
| oSS7 | GGTTAAAGCTTTGGTAGTGCAGGTAATTCG | R Q5 7GC (flank 2)-TAA (pSS3) |
| oSW1 | GCCCCCCGGGACCTGCAGGCAT | F Q5 7GC (flank 1) (pSW2x) |
| oSW2 | GGCCCTGCAGACATGGGTGATCCTCATG | R Q5 7GC (flank 1) (pSW2x) |
| oSW3 | GGGCGGCCAGCCCGCCTAATGAGCGG | F Q5 7GC (flank 2)-TAA (pSW2, pSW3) |
| oSW4 | CCCGGTTAAAGCTTGCATGCCTGCAGG | R Q5 7GC (flank 2)-TAA (pSW2) |
| oSW5 | GCCCCCCGGGATACGCACAATA | F Q5 7GC (flank 1) pSW3x |
| oSW6 | GGCCCTGCAGACATGGGTGATC | R Q5 7GC (flank 1) (pSW3x) |
| oSW7 | CCCGGTTAAAGCTTTGGTAGTGCAGGTAATTCG | R Q5 7GC (flank 2)-TAA (pSW3) |

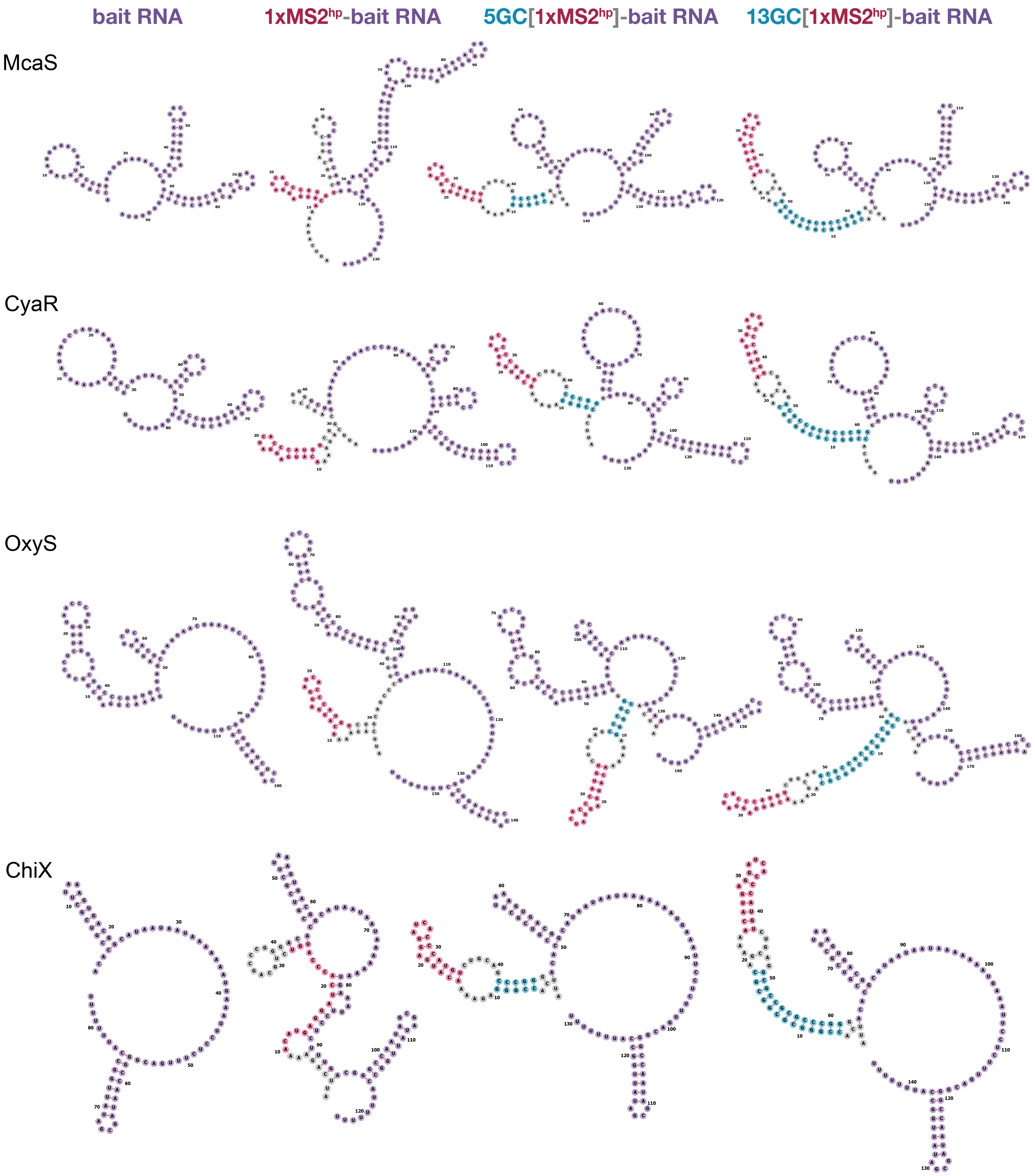


**Supplemental Figure S1. Predicted secondary structures for sRNA pBait constructs.** Secondary structure predictions of *E. coli* sRNAs used in this study (McaS, CyaR, OxyS and ChiX) as isolated sRNA sequences (column 1) and the corresponding hybrid RNAs when each sRNA is inserted into the 1xMS2$^{hp}$ pBait plasmid (column 2) or the short or long GC-clamp constructs (5GC[1xMS2$^{hp}$] and 13GC[1xMS2$^{hp}$]; column 3 and 4, respectively). This panel of sRNA pBait constructs corresponds to data in Fig. 2. RNA structure predictions and visualizations here and throughout the paper were generated using forna (Kerpedjiev et al. 2015).

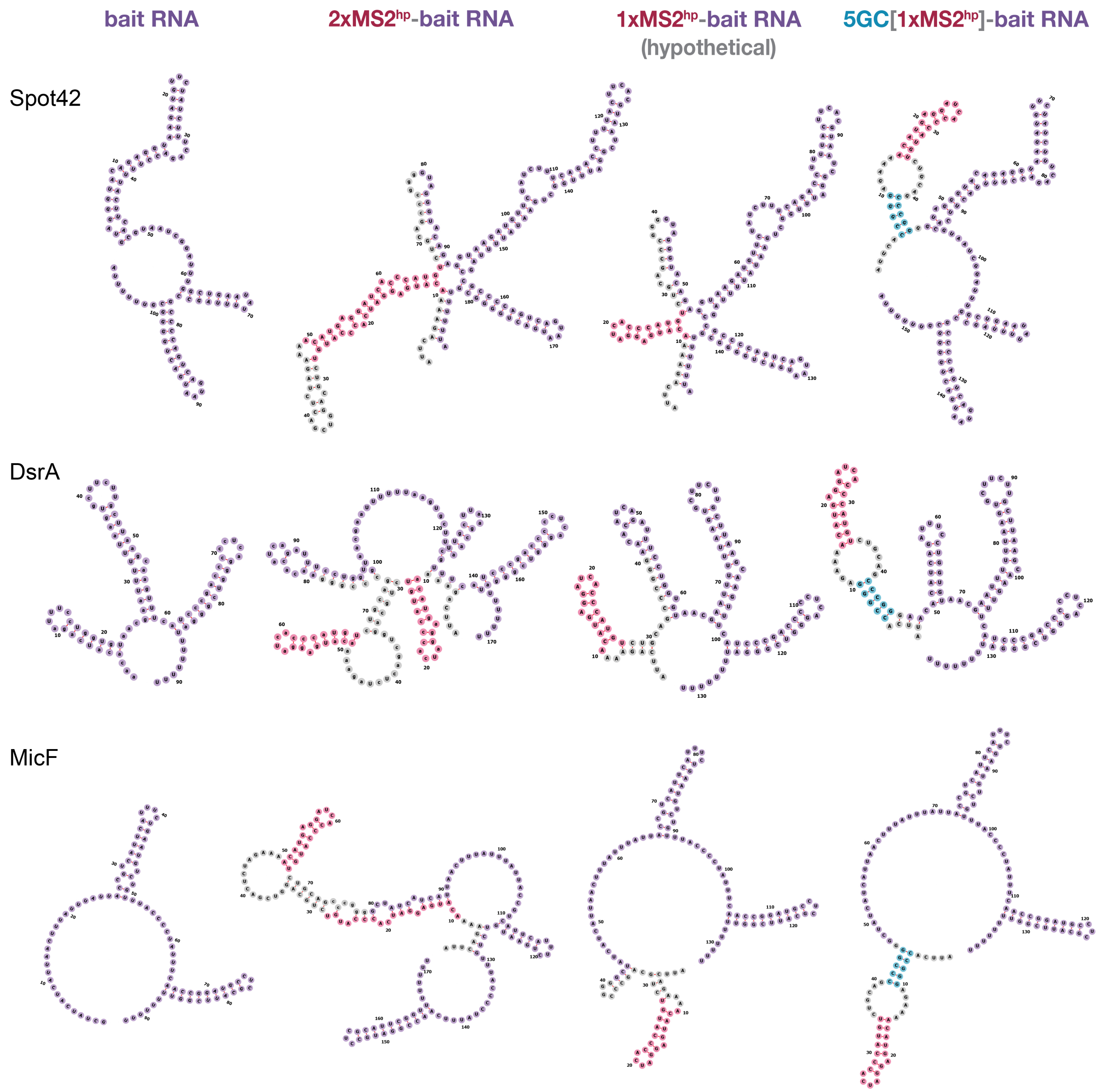


**Supplemental Figure S2. Predicted structures of sRNAs originally cloned into 2xMS2$^{hp}$ pBait constructs (group 1 of 2).** Secondary structure predictions of *E. coli* Spot42, DsrA and MicF sRNAs on their own (column 1) and the corresponding hybrid RNAs when each bait RNA was inserted into the original 2xMS2$^{hp}$ plasmid (column 2) and the short GC clamp construct (5GC[1xMS2$^{hp}$]; column 4). These pBait constructs correspond to data shown in Fig. 3. For comparison, column 3 shows predictions for the structures of hypothetical hybrid RNAs that would have been made with the 1xMS2$^{hp}$ pBait plasmid, even though, for this set of sRNAs, these constructs have not been cloned or tested in the B3H assay; additional constructs in this set are shown in Supplemental Fig. S3.

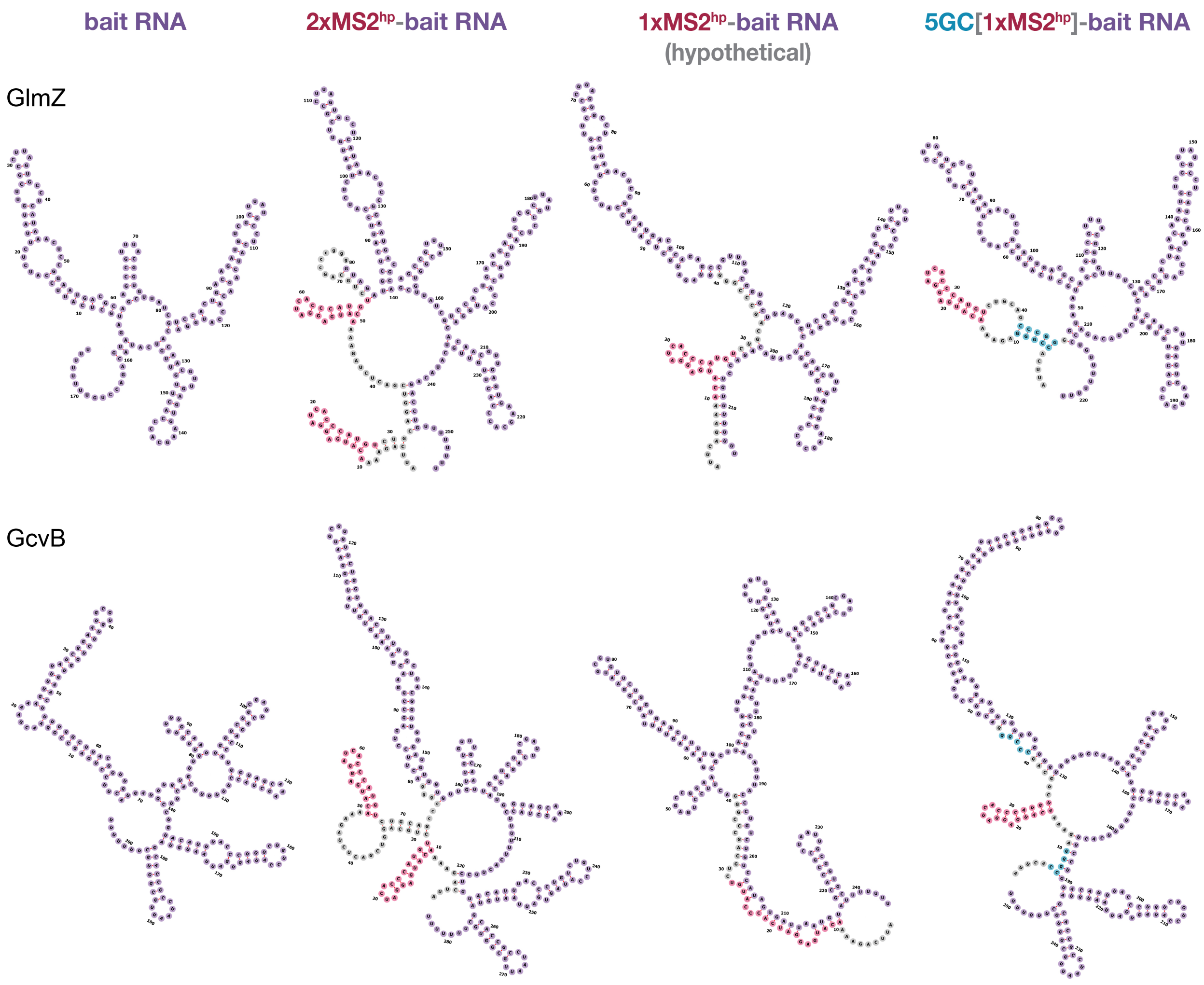


**Supplemental Figure S3. Predicted structures of sRNAs originally cloned into 2xMS2$^{hp}$ pBait constructs (group 2 of 2).** Secondary structure predictions of *E. coli* GcvB and GlmZ sRNAs on their own (column 1) and the corresponding hybrid RNAs when each sRNA was inserted into the original 2xMS2$^{hp}$ plasmid (column 2) and the short GC clamp construct (5GC[1xMS2$^{hp}$]; column 4). These sRNA pBait plasmids correspond to data shown in Fig. 3. For comparison, column 3 shows predictions for the structures of hypothetical hybrid RNAs that would have been made with the 1xMS2$^{hp}$ pBait plasmid, even though these 1xMS2$^{hp}$ constructs have not been cloned or tested in the B3H assay for this set of sRNAs; additional constructs in this set are shown in Supplemental Fig. S2.

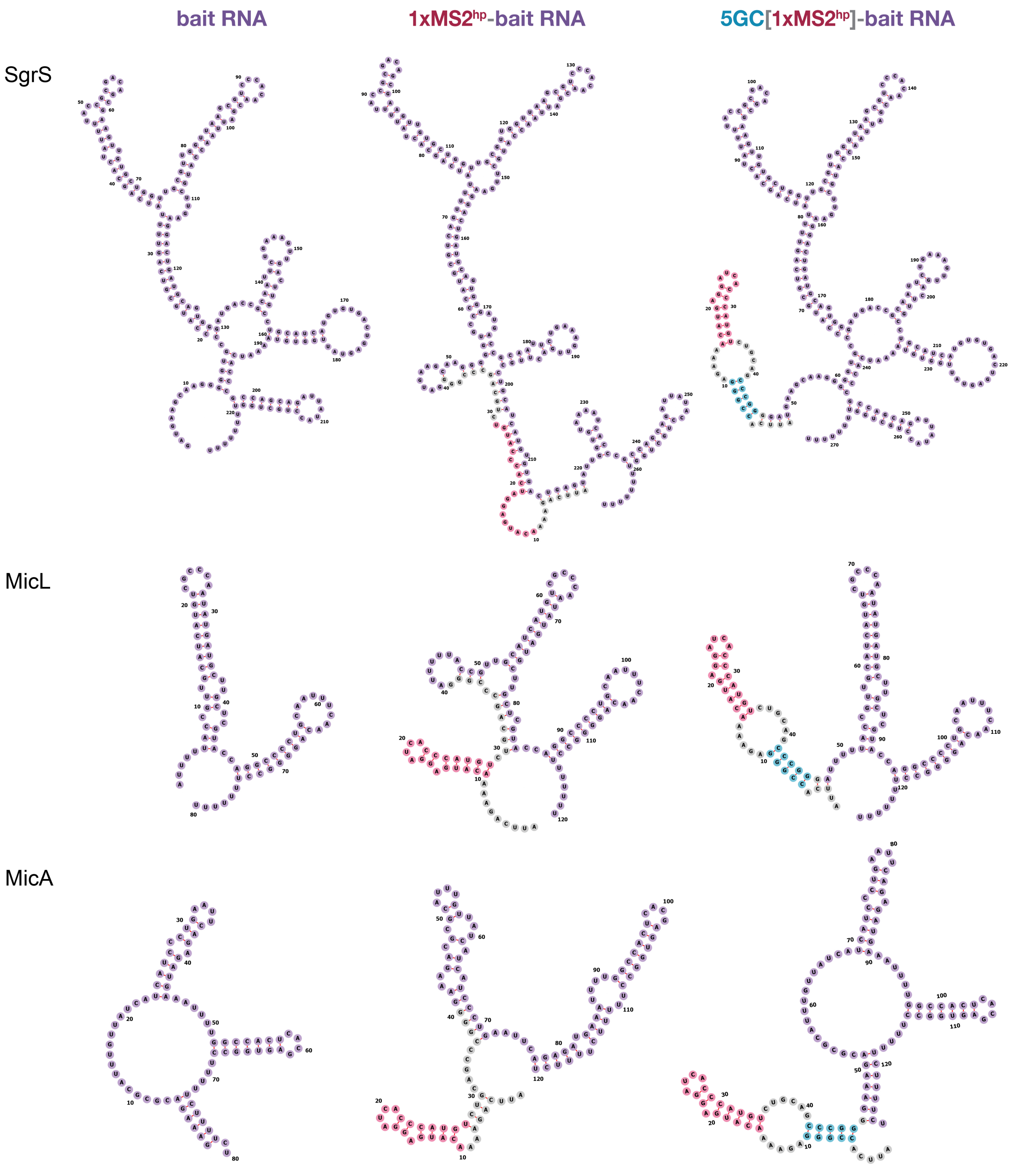


**Supplemental Figure S4. Predicted secondary structures for sRNAs originally cloned into 1xMS2$^{hp}$ pBait constructs and predicted to fold properly in the short GC-clamp constructs (group 1 of 2).** Secondary structure predictions of *E. coli* SgrS, MicL and MicA sRNAs on their own (column 1) and the corresponding hybrid RNAs when each sRNA was inserted into the 1xMS2$^{hp}$ pBait plasmid (column 2) or the short GC-clamp construct (5GC[1xMS2$^{hp}$]; column 3). These constructs represent the set in Fig. 3 that were originally cloned into 1xMS2$^{hp}$ pBait constructs and for which forna predicts that both the sRNA and MS2$^{hp}$ moiety fold properly in the presence, but not the absence, of the 5GC clamp; additional constructs in this set are shown in Supplemental Fig. S5.

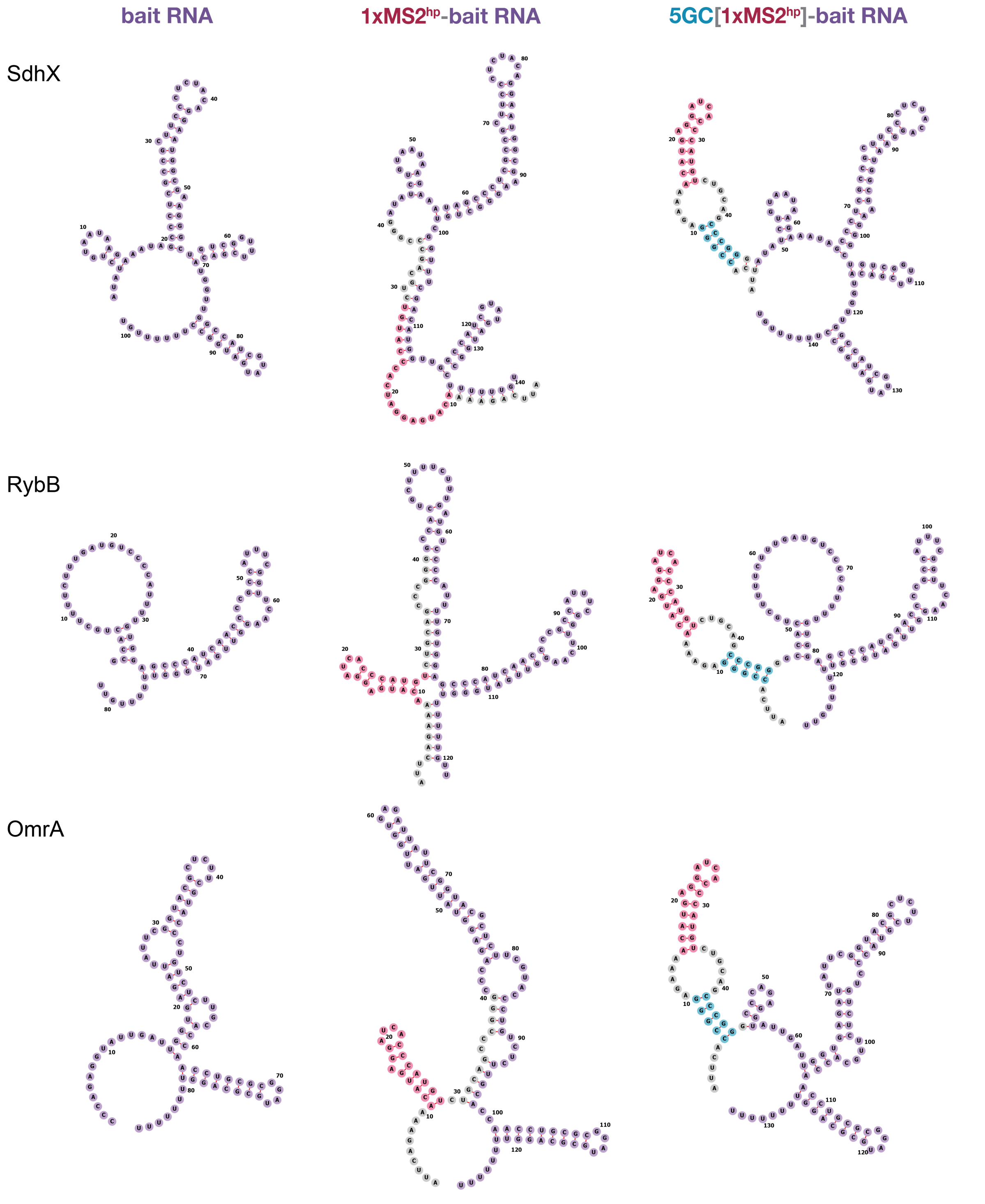


**Supplemental Figure S5. Predicted secondary structures for sRNAs originally cloned into 1xMS2$^{hp}$ pBait constructs and predicted to fold properly in the short GC-clamp constructs (group 2 of 2).** Secondary structure predictions of *E. coli* SdhX, RybB and OmrA sRNAs on their own (column 1) and the corresponding hybrid RNAs when each sRNA was inserted into the original 1xMS2$^{hp}$ pBait plasmid (column 2) or the short GC-clamp constructs (5GC[1xMS2$^{hp}$]; column 3). These constructs represent the set in Fig. 3 that were originally cloned into 1xMS2$^{hp}$ pBait constructs and for which forna predicts that both the sRNA and MS2$^{hp}$ moiety fold properly in the presence, but not the absence, of the 5GC clamp; additional constructs in this set are shown in Supplemental Fig. S4.

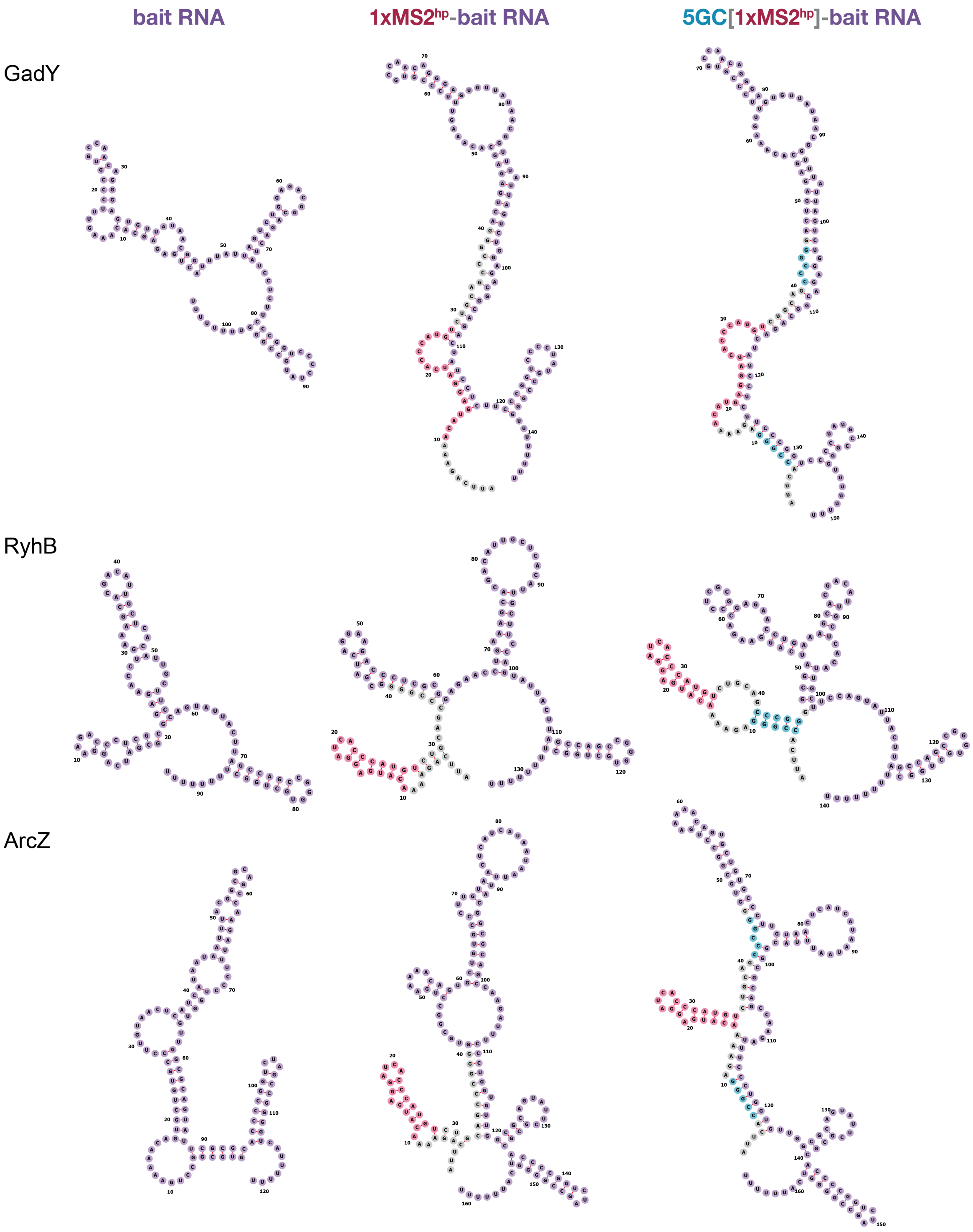


**Supplemental Figure S6. Predicted secondary structures for sRNAs originally cloned into 1xMS2$^{hp}$ pBait constructs and predicted to fold incorrectly in the short GC-clamp constructs.** Secondary structure predictions of *E. coli* GadY, RyhB and ArcZ sRNAs on their own (column 1) and the corresponding hybrid RNAs when each sRNA was inserted into the original 1xMS2$^{hp}$ pBait plasmid (column 2) or the short GC-clamp constructs (5GC[1xMS2$^{hp}$]; column 3). These constructs represent the set in Fig. 3 that were originally cloned into 1xMS2$^{hp}$ pBait constructs and for which forna predicts that either the sRNA and/or MS2$^{hp}$ moiety fold incorrectly in both the presence and absence of the 5GC clamp.

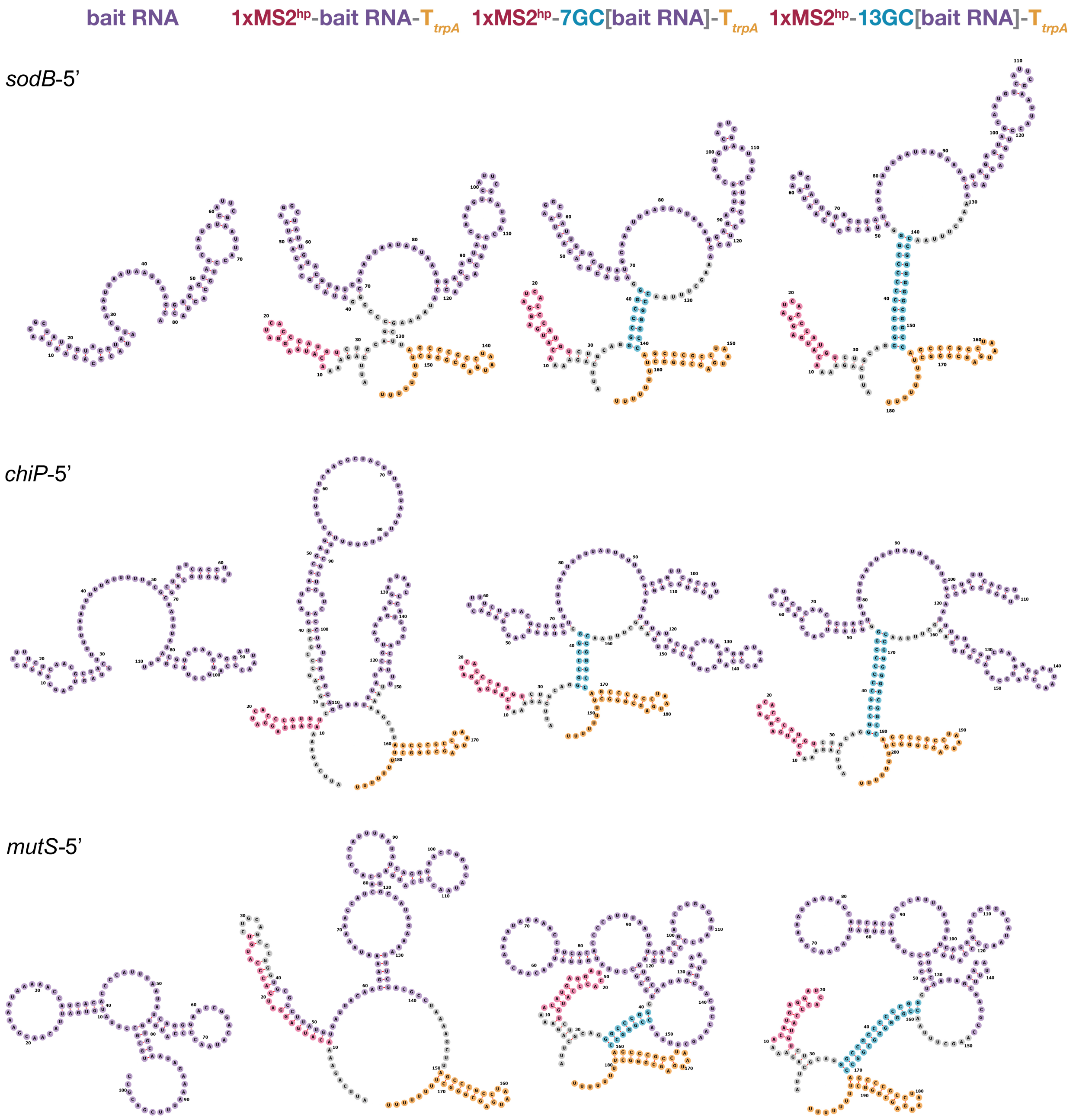


**Supplemental Figure S7. Predicted secondary structures for 5′UTR pBait constructs (group 1 of 2).** Secondary structure predictions of *E. coli sodB*-5′, *chiP*-5′, and *mutS*-5′ UTRs on their own (column 1) and the corresponding hybrid RNAs when each 5′UTR was inserted into the original 1xMS2hp-T*trpA* plasmid (column 2) or the short or long GC-clamp constructs (7GC[bait RNA] and 13GC[bait RNA]; column 3 and 4, respectively). These 5′UTR pBait plasmids correspond to data shown in Fig. 4; additional constructs in this set are shown in Supplemental Fig. S8.

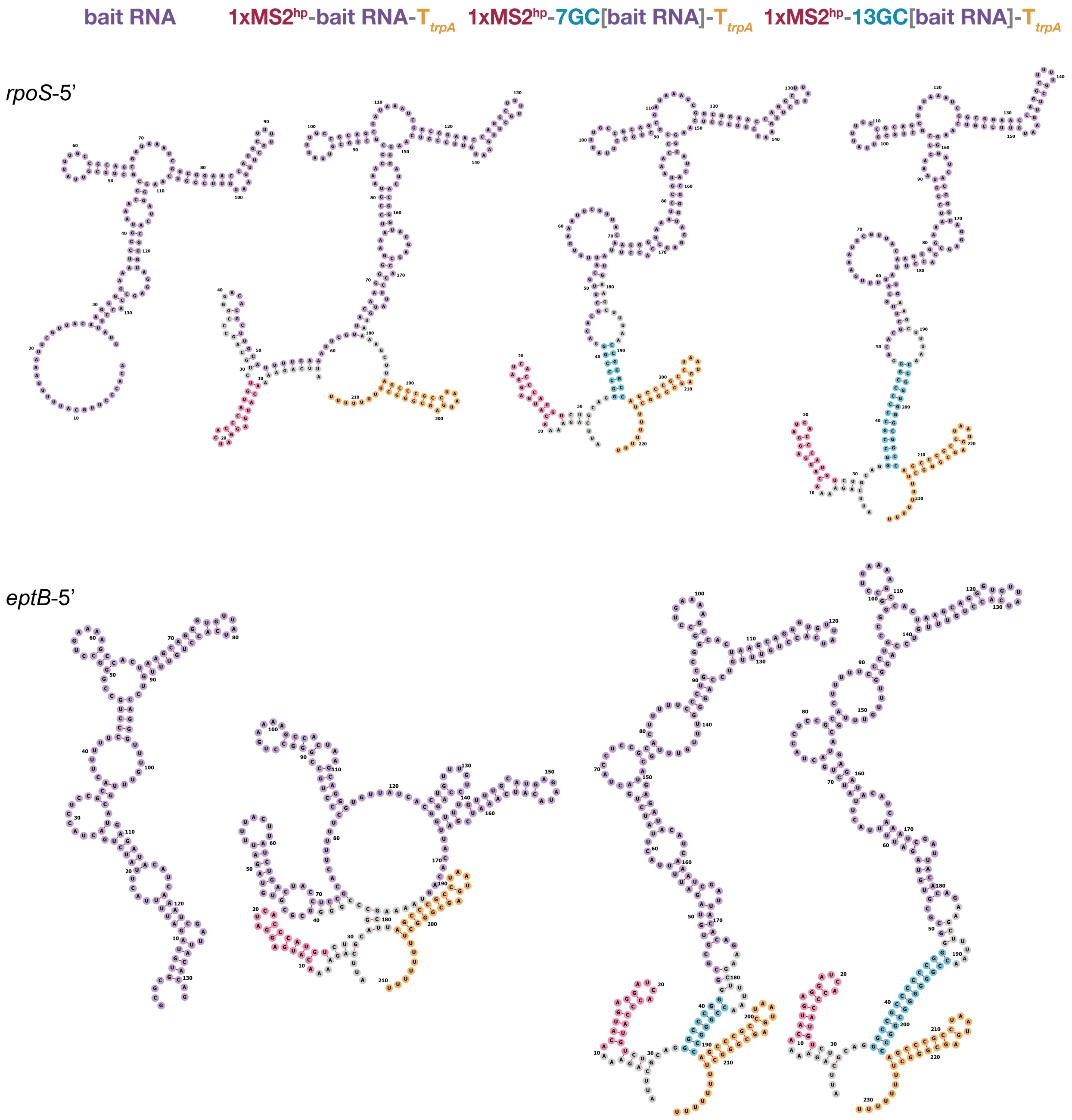


**Supplemental Figure S8. Predicted secondary structures for 5′UTR pBait constructs (group 2 of 2).** Secondary structure predictions of *E. coli rpoS*-5′ and *eptB*-5′ UTRs on their own (column 1) and the corresponding hybrid RNAs when each 5′UTR was inserted into the original 1xMS2$^{hp}$-T*trpA* plasmid (column 2) or the short or long GC-clamp constructs (7GC[bait RNA] and 13GC[bait RNA]; column 3 and 4, respectively). These 5′UTR pBait plasmids correspond to data shown in Fig. 4; additional constructs in this set are shown in Supplemental Fig. S7.
